# Atypical chemoreceptor arrays accommodate high membrane curvature

**DOI:** 10.1101/2020.03.11.986844

**Authors:** Alise R. Muok, Davi R. Ortega, Kurni Kurniyati, Wen Yang, Adam Sidi Mabrouk, Chunhao Li, Brian R. Crane, Ariane Briegel

## Abstract

The prokaryotic chemotaxis system is arguably the best-understood signaling pathway in biology, but most insights have been obtained from only a few model organisms and many studies have relied on artificial systems that alter membrane curvature^1–3^. In all previously described species, chemoreceptors organize with the histidine kinase (CheA) and coupling protein (CheW) into a hexagonal (P6 symmetry) extended array that is considered universal among archaea and bacteria^4,5^. Here, for the first time, we report an alternative symmetry (P2) of the chemotaxis apparatus that emerges from a strict linear organization of CheA in *Treponema denticola* cells, which possesses arrays with the highest native curvature investigated thus far. Using cryo-ET, we reveal that the *Td* chemoreceptor arrays assume a truly unusual arrangement of the supra-molecular protein assembly that has likely evolved to accommodate the high membrane curvature. The arrays have several additional atypical features, such as an extended dimerization domain of CheA and a variant CheW-CheR-like fusion protein that is critical for maintaining an ordered chemosensory apparatus in an extremely curved cell. Furthermore, the previously characterized *Td* oxygen sensor ODP influences array integrity and its loss substantially orders CheA. These results suggest a greater diversity of the chemotaxis signaling system than previously thought and demonstrate the importance of examining transmembrane systems *in vivo* to retain native membrane curvature.

## Introduction

Chemotaxis is a behavior most motile bacteria employ to sense their chemical environment and navigate toward favorable conditions. The main components of the system are transmembrane chemotaxis receptors called <u>m</u>ethyl-accepting <u>c</u>hemotaxis <u>p</u>roteins (MCPs), the histidine kinase CheA, and the adaptor protein CheW. The intracellular tips of MCPs bind CheA and CheW and communicate changes from the external chemical environment into the cell by modulating CheA kinase activity (Fig. S1A)^6–8^. Activation of CheA initiates an intracellular phosphorelay that ultimately controls flagellar rotation and cell movement. CheA functions as a dimer and possesses five domains (P1-P5) with distinct roles in autophosphorylation and array integration. The P1 domain is the phosphate substrate domain, P2 interacts with response regulators, P3 is the dimerization domain, P4 binds ATP, and P5 interacts with CheW. In the model species *Escherichia coli (Ec),* CheA P5 and CheW are paralogs that interact pseudo-symmetrically to form six-subunit rings. In all bacterial and archaeal species examined thus far, the MCPs are arranged in a trimer-of-dimer oligomeric state and further organize into a hexagonal lattice (Fig. S1B). In *Ec,* the receptors are connected by the highly ordered rings of CheA and CheW bound to the cytoplasmic tips of the receptors^5,9^. These insights have established a widely accepted central model of the chemotaxis array (Fig. S1B)^4,5,10^. However, emerging research has recently revealed divergent components and arrangements of the chemotaxis apparatus in non-canonical organisms. For example, in *Vibrio cholerae (Vc)* chemotaxis arrays, CheA and CheW lack an ordered arrangement in the rings^1,11^.

Many of these structural insights have transpired from cryo-electron tomography (cryo-ET) studies that utilize artificial systems for higher resolution data. Specifically, the advent of so-called ‘mini-cell’ bacterial strains produce extremely small cells that are ideal for cryo-ET^1,3^, and lipid-templating methods generate *in vitro* arrays with increased conformational homogeneity^2,12^. However, these methods generate arrays with non-native curvature and it is unclear how this may affect array structure and behavior. Here, we examine the *in vivo* array structure of the pathogenic spirochete *Treponema denticola (Td)* by cryo-ET, which possesses the highest native membrane curvature of any bacteria examined for their chemotaxis system so far^4,5^. We demonstrate the presence of a novel array architecture with two-fold (P2) symmetry in *Td*, that likely owes to the high curvature of the cells. Genetics experiments, bioinformatics analyses, structural investigations, and molecular modeling of *Td* chemotaxis proteins reveals adaptations that have likely evolved to accommodate formation of an extended chemotaxis array in a highly curved membrane. We demonstrate that a CheR-like fusion domain in a variant CheW is key for maintaining the structural integrity of the arrays. Furthermore, cryo-ET analysis of *Td* cells lacking the oxygen sensor ODP reveals substantial changes to the ordering or mobility of CheA^13^. Collectively, these data demonstrate a greater diversity of the chemotaxis system than previously assumed, and exemplify the importance of examining biological structures in native *in vivo* conditions.

## Results

### Conservation of the F2 chemotaxis system

The chemotaxis systems in prokaryotes have been classified into 19 systems based on phylogenomic markers^14^. These classes include 17 ‘<u>F</u>lagellar’ systems (F), one ‘<u>A</u>lternative <u>C</u>ellular <u>F</u>unction’ system (ACF), and one ‘<u>T</u>ype <u>F</u>our <u>P</u>ilus system’ (TFP). The spirochete chemotaxis system belongs to the ‘Flagella class 2’ (F2) category, which has not been investigated with structural methods^14^. To explore the characteristics of chemotaxis proteins in F2 genomes, we analyzed genomes in the Microbial Signal Transduction Database version 3 (MiST3)^15^. All genomes (306) with at least one CheA of the F2 class (CheA-F2) are in the Spirochaetota phylum, with a few exceptions of lateral gene transfer to other phyla (Dataset 1, see Supplementary Methods). Not all Spirochaetota bacteria possess an F2 system however. There are 1096 genomes assigned to the Spirochaetota phylum in GTDB^16^ and 804 of these are present in the MiST3 database (Dataset 2)^15^. With these 804 genomes, we mapped the different classes of CheA kinases to the Spirochaetota taxonomy tree (Figure S1). Based on the topology of this tree, it appears that the major chemosensory systems in the genomes from the Spirochaetota phylum are: F1 (Leptospirae), a transitional hybrid F1/F2 system (Brachyspirae), and F2 (Spirochaetia) (Fig. S2). Therefore, we conclude that F2 systems are exclusive to the Spirochaetia class, with a few exceptions of lateral gene transfer.

The main architectural difference of the F2 system compared to others is the presence of an unusual scaffold protein that consists of an N-terminal CheW domain and a C-terminal CheR-like domain, hereafter referred to as CheW-CheR_like_. Typically, CheR is a methyltransferase that, together with the methylesterase CheB, controls the methylation state of the receptors and thus provides an adaptation system^17^. Our analyses indicate that all Spirochaetia genomes in MiST3 contain CheW-CheR_like_. Furthermore, if we limit our analysis to genomes that are fully sequenced (see Supplementary Methods), CheW-CheR_like_ is only found in F2 chemosensory systems.

To investigate sequence patterns in the CheR protein and the CheR-like domain of CheW-CheR_like_, we produced a sequence dataset with 83 CheR-F2 and 88 CheW-CheR_like_ proteins and summarized them in sequence logos (Fig. S3A). Although the key catalytic residues are conserved in the typical CheR protein, two residues that are essential for CheR to methylate chemoreceptors, R79 and Y218 in *Td* CheR, are modified in the CheW-CheR_like_ protein (R79W and Y218F) ^17^. Furthermore, the conserved region at the C-terminus of CheR is not conserved in CheW-CheR_like_. Based on these results we speculate that the CheR_like_ domain does not possess methyltransferase activity. Collectively, our analyses suggest that CheR and CheW-CheR_like_ have different biological functions.

F2 systems contain three proteins with a CheW domain: the classical scaffold CheW, CheW-CheR_like_, and the P5 domain from the histidine kinase CheA. Phylogenetic mapping of CheW-F2 proteins indicate the location of last common ancestor (Fig. S4, see Supplementary methods). To investigate sequence patterns of the three CheW domains, we analyzed non-redundant sequence datasets of CheW, CheW-CheR_like_, and CheA-F2 from all genomes with at least one CheA-F2. The final alignments for each group contain the CheW domain portion of 74 CheW proteins, 59 CheW-CheR_like_ proteins, and 73 CheA proteins. The sequences of each group are summarized in sequence logos and demonstrate the presence of conserved regions at established interaction interfaces, as well as loop insertions near these interfaces that could confer altered specificity of binding (Fig. S3B).

### The structure of the *Treponema denticola* (*Td*) chemotaxis array in wild-type cells

Cell poles of intact *Td* cells were imaged by cryo-electron tomography (cryo-ET) and used for three-dimensional reconstructions. Top views (cross-sections through the array) and side views (visualizing the long axes of the receptors) of membrane-associated arrays were clearly visible (Fig. 1A, S5A). Sub-volume averaging revealed the conserved receptor trimer-of-dimers in the typical hexagonal arrangement. Remarkably, several novel features of the chemotaxis arrays are apparent (Fig. 1B). Specifically, a density of unknown origin is located in the center of the receptor hexagons and slightly above the plane of the CheA:CheW rings. This density, which will hereafter be referred to as the middle density, extends from two subunits in the rings (Fig. 2A). Additionally, there are small but distinct puncta of density in between some of the trimer-of-dimer modules (Fig. 1B). However, averages of the arrays at the CheA:CheW layer did not reveal discernible CheA density, indicating either a sparse or disordered distribution of CheA or a highly mobile kinase (Fig. 1C). Remarkably, the subtomogram averages reveal the axis of the *Td* cells relative to the chemotaxis arrays, demonstrating that the arrays occupy a preferred orientation with respect to the cell axis (Fig. S5B).

**Fig. 1.**
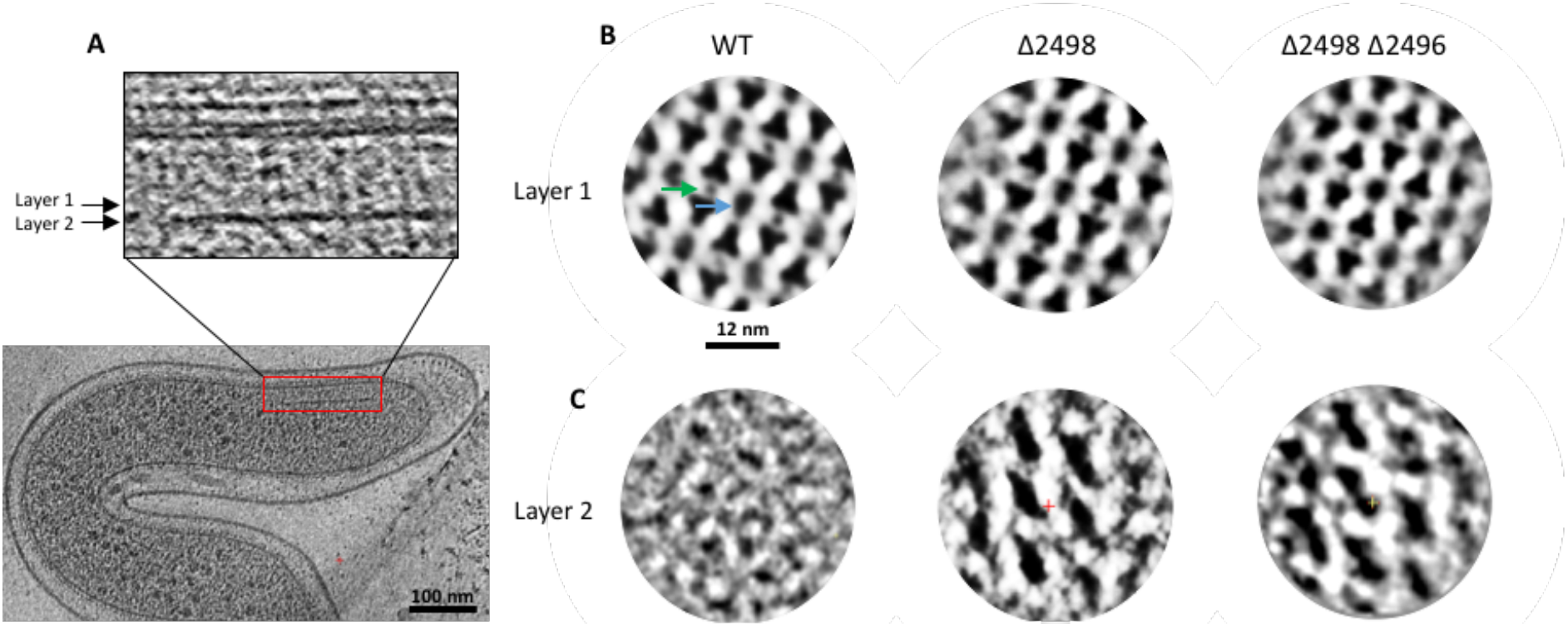
Cryo-electron tomography of whole *T. denticola* (*Td*) cells reveals the protein arrangement of chemotaxis machinery. (A) Side-views of the membrane-associated chemotaxis apparatus illustrate the location of the receptor layer (Layer 1) and CheA:CheW baseplate (Layer 2). These layers are spaced ∼90 Å from one another. (B) Sub-volume averaging of three *Td* strains reveals the universally-conserved receptor trimer-of-dimer arrangement with 12 nm spacing between opposing trimer-of-dimer modules. Notably, density is apparent in the center of the receptor hexagons (blue arrow) and between some receptor trimer-of-dimer modules (green arrow). (C) Sub-volume averages at Layer 2 reveal the organization of CheA. In this arrangement, the density between the trimer-of-dimer modules (green arrow) corresponds to the CheA P3 domain.

**Fig. 2.**
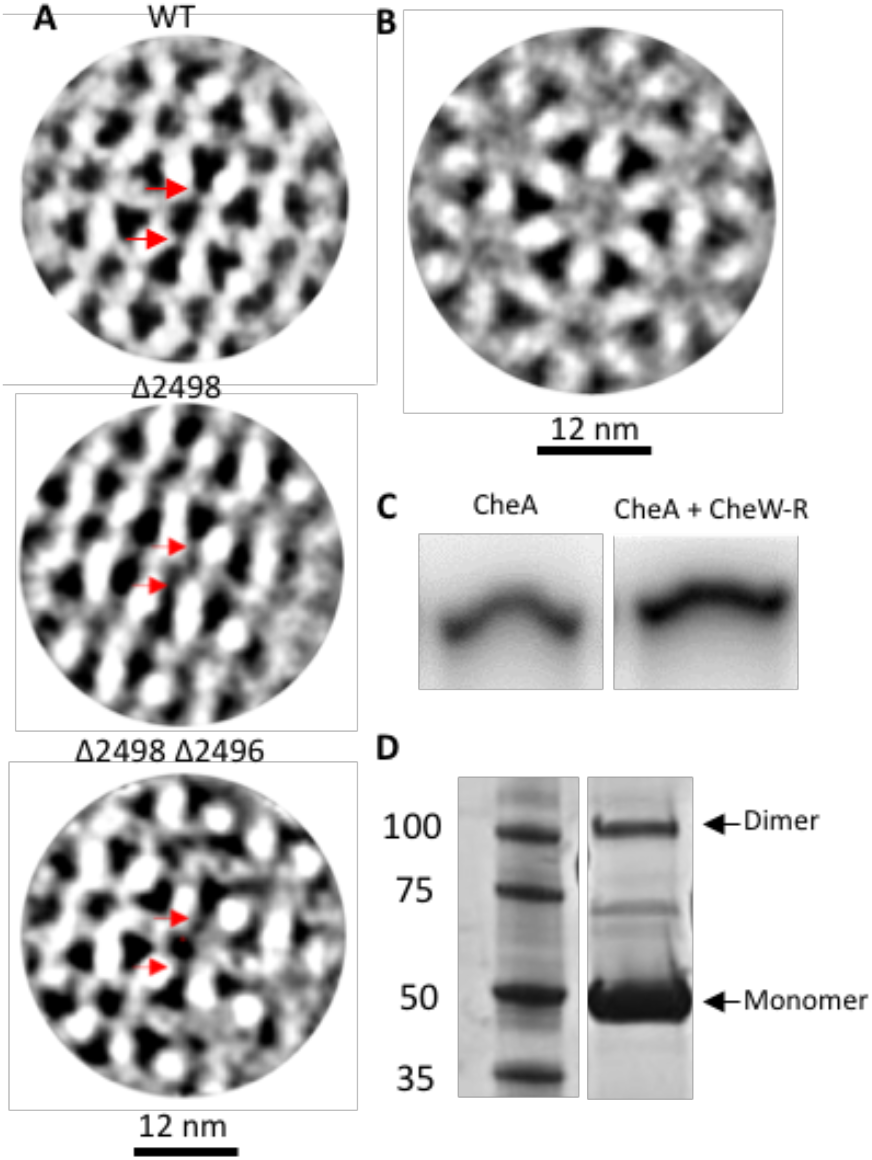
Investigations into the identity of the density located in the center of the receptor hexagons, referred to as the middle density. (A) In Layer 1, the middle density extends to two positions in the CheA:CheW rings for all three *Td* strains (red arrows). (B) Deletion of a CheR-like domain that is covalently attached to the C-terminus of a CheW domain generates sub-tomogram averages that lack the middle density feature. (C) Radioisotope assays that monitor CheA autophosphorylation with [γ-^32^P]ATP demonstrate that CheW-CheR_like_ influences CheA activity *in vitro.* Bands are from non-adjacent lanes on the same gel. (D) Purification of CheW-CheR_like_ followed by Native PAGE gel analysis reveals the presence of a protein monomer (49 kDa) and protein dimer (98 kDa). Bands are from non-adjacent lanes on the same gel.

### Arrays in *T. denticola* deletion mutants

The <u>o</u>xygen-binding <u>d</u>iiron <u>p</u>rotein (ODP) functions as an oxygen sensor for chemotaxis in *Td*, however it is uknown if ODP is an integral component of the array^13^. This protein is genetically coupled to an MCP homolog, TDE2496, that lacks both transmembrane and sensing modules. TDE2496 likely integrates into the cytoplasmic regions of the membranebound arrays based on the observation that no cytoplasmic arrays were observed in the tomograms. Moreover, the *Td* genome encodes only one CheA homolog, and cytoplasmic receptors often associate with distinct kinases^9,18^. TDE2496 forms trimers-of-dimers *in vitro* and can modulate CheA activity (Fig. S6A,B). To determine if the presence of ODP (TDE2498) and its cognate receptor (TDE2496) impacts array architecture or integrity, we conducted cryo-ET with two *Td* gene knock-out strains, Δ2498 and Δ2498Δ2496^13^. Importantly, deletion of ODP does not impact transcription of TDE2496^13^. The sub-volume averages of these strains reveal distinct differences in array densities compared to the wildtype (WT) strain (Fig. 1B,C, Fig. S5A). Namely, the location of CheA at the CheA:CheW layer (Layer 2, Fig 1C) is now clearly visible. Interestingly, CheA arranges in well-ordered linear rows. Placement of CheA necessarily positions the P3 domain in between two of the trimer-of-dimer modules in each hexagon. This position exclusively corresponds to the location of the puncta between receptor trimer-of-dimer modules observed in the WT array, indicating that this density represents the P3 domain (Layer 1, Fig. 1B). Like the WT arrays, the cell axis relative to the chemotaxis arrays is also apparent in these sub-tomogram averages and matches the preferred array orientation in the WT cells (Fig. S5B).

### Analyses of the CheW-CheR_like_ protein in *T. denticola*

To determine the composition of the middle density in the *Td* arrays, we considered the unique chemotaxis proteins in spirochete genomes. As shown above, CheW-CheR_like_ is a conserved component of the F2 Spirochaetia chemotaxis system (Fig. S2). In *Td*, a 28-residue linker in CheW-CheR_like_ separates the two domains and is predicted to be largely helical (JPred), and the gene is co-transcribed with the only CheA, CheX, and CheY proteins in the genome (Fig. S6C,D)^18^. Furthermore, the purified CheW-CheR_like_ protein dimerizes and interacts with CheA, as evidenced by its ability to alter autophosphorylation activity (Fig. 2C,D, Fig. S6E,F). Therefore, we postulated that the middle density may be comprised of two dimerized CheR_like_ domains that extend from the CheW units harbored within the CheA:CheW rings.

A *Td* strain lacking the CheR_like_ domain (ΔCheR_like_) reveals a significant decrease in the prevalence and size of the arrays (Fig. S5A). Due to the small size of the arrays in ΔCheR_like_, only 194 particles were available for sub-tomogram averaging, but the resulting averages clearly demonstrate that the middle density is no longer present (Fig. 2B, S5A). Like the WT strain, the CheA density below the rings in ΔCheR_like_ is not apparent.

### Protein interfaces in the CheA:CheW:CheW-CheR_like_ rings

Bioinformatics analyses demonstrate that all functional Spirochaetota F2 chemotaxis systems possess a CheW-CheR_like_ homolog and at least one classical CheW protein. As only two of the CheW subunits in the hexagonal rings extend to the middle density (which probably arises from the CheW-CheR_like_ protein), and two of the ring positions are occupied by CheA P5, it is likely that the other two positions are occupied by the classical CheW protein. (Fig. 2A, S7). Within this arrangement three unique interaction interfaces are possible, interface 1 occurs between CheA P5 and the classical CheW (as seen in canonical systems), interface 2 occurs between CheA P5 and the CheW domain of CheW-CheR_like_, and a third interface (interface 3) occurs between the classical CheW and the CheW domain of CheW-CheR_like_ (Fig. S7). To explore the binding interfaces within the *Td* rings, we analyzed homology models of the classical CheW, the CheW domain of CheW-CheR_like_, and the CheA P5 domain. The CheW models were generated using a crystal structure of *Thermoanaerobacter tengcongensis* (*Tt*) CheW as the template (PDB ID: 2QDL), and the CheA P5 model was generated using a cryo-EM structure of *E. coli* CheA P5 (PDB ID:6S1K) (Fig. S8A-C)^2,19^. Three of the four regions with lowest sequence conservation among the three domains are located at interfaces 1-3 (Fig. S9). Alignment of the *Td* CheW and CheA P5 models to a crystal structure of *Tm* CheW in complex with *Tm* CheA P5 (PDB ID: 3UR1) further illustrates that these regions are located at the CheW:P5 ring interfaces (Fig S9)^20^. Mapping the variable regions onto the sequence logos of the F2 CheW domains demonstrates that they evolved different sequence patterns in these regions, with the exception of the variable region that is not located at the interaction interface (region 2, Fig. S9, S3B).

### CheA arrangement and array curvature in *T. denticola*

Sub-tomogram averaging reveals that *Td* CheA forms a linear arrangement across the chemotaxis array, linking the CheA:CheW rings into extended ‘strands’ that are held together by receptor:CheA/W interactions (Fig. 3A,B). The orientation of the strands are apparent in the sub-tomogram averages and run relatively perpendicular to the axis of the cells (Fig. S10A). The array densities at Layer 1 (receptors, P3, and CheR_like_) produce apparent lines in the cryo-ET reconstructions that, like the sub-tomogram averages, run parallel to the cell axis (Fig. 3C,D)^21^. This arrangement allows interactions that hold the strands together to occur with minimal bending (Fig. 3E,F). Indeed, the angle between the cell axis and these lines in the cells is 10.4 +/− 8.6°, (*n* = 26 cells) and no significant difference was found among the three *Td* strains measured (Table S1A). As the CheA:CheW strands are perpendicular to these lines in the sub-tomogram averages, they will be oriented at an angle of 55.4 +/− 8.6° with respect to the cell axis (Fig. S10A).

**Fig. 3.**
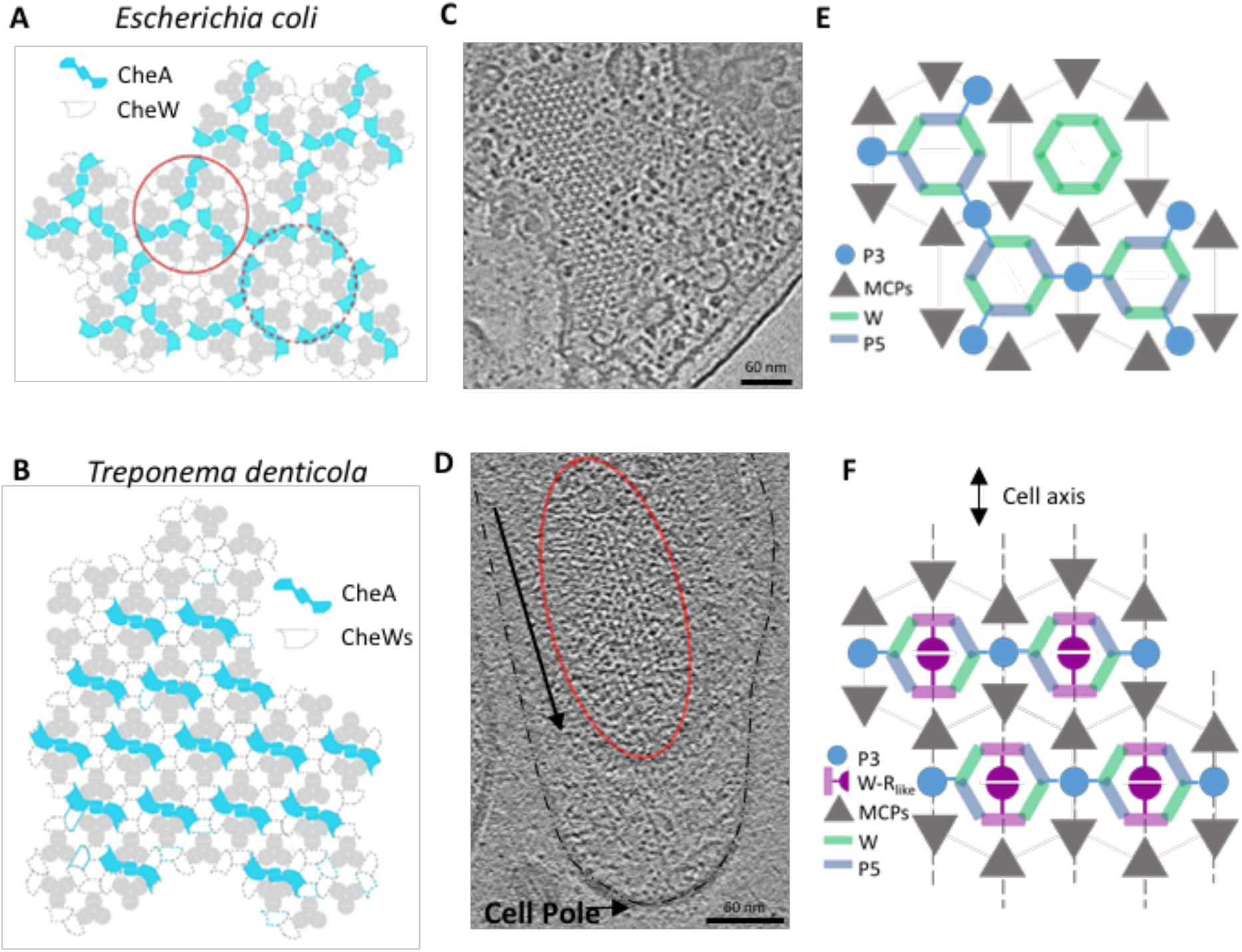
The organization of CheA in *Td* differs from all previously reported arrangements. (A) The hexagonal arrangement of CheA in *E. coli* and other canonical chemotaxis systems. (B) In *Td,* CheA is arranged with a strict linear organization, producing ‘strands’ of CheA:CheW rings linked by the CheA P3 domain. (C) The receptor densities in *Ec* (Layer 1) are apparent and hexagonal. Figure adapted from Briegel A. et al. 2010.^21^ (D) The densities at Layer 1 in *Td* (red oval) produce apparent lines of receptors, P3 and the middle density (CheR-like) that follow the axis of *Td* cells (black arrow). (E) *Ec* arrays possess radial P6 symmetry where all interaction interfaces are equally curved. (F) The *Td* array has a two-fold symmetry arrangement that allows receptor:CheA/W interactions that hold the CheA:CheW strands together to occur with minimal bending. Dashed lines represent the apparent lines in the tomograms.

Previous studies in *Ec* and *Vc* demonstrate that the chemotaxis arrays can accommodate vastly different curvatures between lysed and artificially small mini-cell strains^1,3,22^. However, *Td* cells demonstrate a significantly higher curvature of the cell membrane than organisms investigated for their chemotaxis arrangement thus far^1,5,22^. As a measure of comparison, the *Vc* mini-cells used in a previous study have an inner membrane curvature of 9.15 +/− 4.5 /μm (radius 1092 Å, *n* = 6 cells), and the *Td* cells have an inner membrane curvature of 35.8 +/− 6.6 /μm (radius: 279 Å, *n* = 10 cells) (Fig. S10B,C, Table S1B,C). Additionally, the measured curvature of the *Td* CheA:CheW baseplate is 65.6 +/− 19 /μm (radius: 152 Å, *n* = 10 cells) (Fig. S10B, Table S1C).

The CheA:CheW rings present in a crystal structure (PDB ID: 3UR1) are flat and ∼95 Å in diameter (Fig. S11A)^20^. The length across two rings connected by a dimeric CheA is 224 Å (Fig. 11B). To determine the extent of bending that would need to occur in the two CheA:CheW rings if they ran perpendicular to the cell axis, the 224 Å rings were modeled as a chord in a circle with radius 152 Å. Using the equation 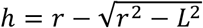 (where h is the height of the circular segment, r is the circle radius, and L is half the chord length (224 Å /2)), the height of the circular segment is 49.2 Å (Fig. S11B). Therefore, the center of the two rings (the P3 domain) must bend by an average of 49.2 Å toward the cell membrane to accommodate the cell curvature. Using the same equation above (where L is 95 Å /2), the height of the circular segment is 7.6 Å (Fig. S11A). Therefore, in *Td,* the center of a single ring must bend toward the membrane by an average of 7.6 Å to align to the measured baseplate curvature.

### Spirochetes possess an atypical dimerization domain

The cryo-ET results reveal density corresponding to the P3 domain, which has not been previously reported in *in vivo* arrays. Sequence alignments of *Td* CheA with CheA homologs from a variety of model bacteria with previously characterized chemotaxis systems indicate that in *Td* CheA an additional ∼50 residues join the canonical dimerization domain (P3) helices (Figure S12A)^4^. CheA homologs from other spirochete genera including Borrelia and Brachyspira also possess additional residues in this region (Fig. S12B)^23^. Analysis of non-redundant P3 domains from all CheA classes reveal general sequence conservation in the canonical helices but highly divergent sequences at these additional residues (Fig. S13A). Furthermore, CheA-F2 proteins contain the most residues in this non-conserved region (Fig. S13B,C). The x-ray crystal structure of the isolated *Td* P3 domain (PDB ID: 6Y1Y, Fig. 4, Table S2) reveals that the additional residues adopt the coil-coiled motif of the classic dimerization domain with the exception of a break in one of the helices, producing a discontinuous coiled-coil (Fig. 4, S14A). Interestingly, aromatic residues (Phe, Tyr) cluster near the helix breakages and unusual core packing of these residues allows for maintenance of a coiled-coil register despite a disruption of helical heptad repetition in the C-terminal helix (Fig. S14A). The net result is a distortion in the alignment of the hairpin tip, the consequence of which is currently unknown. Differing orientations of Tyr83 in the two subunits produces asymmetry in the added tip region (Fig. S14B,C). Fitting the new P3 domain into an all-atom chemotaxis array that was generated for previous molecular dynamics simulations (PDB ID: 3JA6) shows that these additional helices are within a ∼15 Å from receptors (Fig. S14D)^12^. Additionally, the handedness of the helix connection in the *Td* P3 domain differs from that of *Thermotoga maritima* CheA and rather matches helix connectivity of sensor kinase DHP domains^24,25^.

**Fig. 4.**
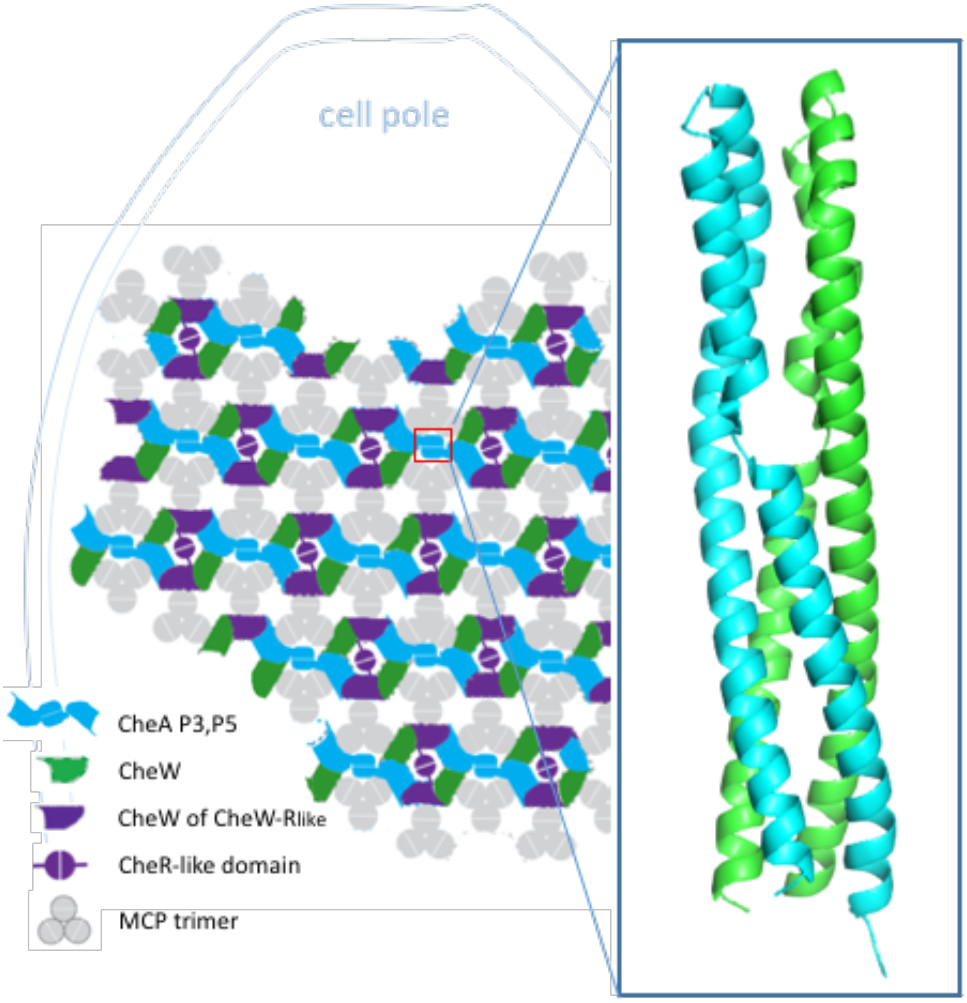
The arrangement of chemotaxis proteins in *Td*. The linear organization of CheA produces ‘strands’ of CheA:CheW rings that are perpendicular to the cell axis and are linked by an atypical P3 domain.

## Discussion

Here, we reveal the protein arrangement of F2 chemotaxis arrays through cryo-ET of intact *T. denticola* (*Td*) cells. *Td* cells are thin and cyndrillical, producing a cell that is polarized in shape and membrane curvature; the membrane has extremely high curvature perpendicular to the cell axis and lower curvature parallel to the axis. In accordance with this feature, the *Td* arrays are polarized and assume a preferred orientation with respect to the cell axis. In this system, three proteins comprise the rings at the receptor tips: CheA, CheW, and a CheW-CheR_like_ protein. Like the *Ec* system, these proteins in *Td* are integrated into the array with strict organization^1,22^. However, a linear arrangement of CheA is present that generates ‘strands’ of rings interlinked by the CheA dimerization domain (P3). The strands run perpendicular to the axis of the *Td* cells, resulting in substantial bending of inter-linked rings. Presumably, the unique binding interfaces in the CheA:CheW rings generate bent interfaces with the appropriate curvature. Furthermore, this directional distinction allows for some array contacts to follow the cell axis and remain undistorted. The deleterious effect on array integrity with the loss of the CheR_like_ domain strongly suggests that CheR_like_ dimerization plays a key role in array assembly and stabilization. The strict linear arrangement of CheA could be facilitated by the composition of the *Td* rings; three unique protein interfaces are present in the rings and restrict CheA P5 integration (i.e. CheA P5 can only occupy these two positions in the six-member ring). Furthermore, the *Td* CheA P3 domain is clearly discernible in the averaged tomograms, which has not been previously observed *in vivo^5,9,22^.* As the CheA:CheW rings have to undergo substantial bending to accommodate the baseplate curvature, the elongated P3 domain may have evolved to stabilize CheA dimerization by increasing the interface area and furthermore may encourage interactions with neighboring receptors^12^. Due to the curvature, the receptor trimer-of-dimer modules are expected to be further splayed, and the elongated P3 may compensate for the increased distance between receptors and P3. As *Td* cells have the smallest average diameter (0.1 – 0.4 μm) of all bacteria with determined chemotaxis architectures thus far, and the novel linear arrangement of CheA produces arrays that can better accommodate its unique membrane curvature, we surmise that the array architecture in *Td* is an adaptation that evolved to produce an extended chemoreceptor apparatus in a highly curved and asymmetric membrane^4,5,9^. Importantly, bioinformatics analyses indicate that the unique protein features seen in *Td* are exclusive to Spirochaetia F2 systems.

Unexpectedly, the placement of CheA in WT *Td* arrays could not be discerned (with the exception of the P3 domain), but was clearly visible in two *Td* mutants (Δ2498, Δ2498Δ2496). As the density corresponding to the P3 domain in the WT strain is clearly discernible, the sparse density corresponding to all other CheA domains (P1, P2, P4, P5) is not attributed to low CheA incorporation in these arrays. These results indicate that the kinase is highly mobile or more disordered in the WT strain, but is more constrained when ODP (TDE2498) is deleted, suggesting that ODP directly affects array structure. However, densities in the three strains do not designate an obvious position for ODP, indicating that ODP may not be an integral component of the array, but rather peripherally interacts with the chemotaxis machinery or influences its architecture through other means, perhaps related to its signaling properties.

In summary, we illustrate a novel chemotaxis arrangement that has evolved to compliment the spirochetes’ high membrane curvature and asymmetry. Therefore, it is likely that the behavior and characteristics of chemoreceptor arrays in general can be influenced by perturbing the shape of the cell membrane. Recent studies with *Ec* ultra-minicells (the smallest mini-cells available to date) produce densities corresponding to portions of chemoreceptors that have not been observed before^3^. However, it is possible that increased membrane curvature limits movement of the transmembrane receptors, resulting in increased receptor localization and resolution. Furthermore, the use of lipid-templating has been extensively used to assemble arrays *in vitro* for cryo-ET and reconstitutes the chemotaxis apparatus in a perfectly flat formation^2,12^. While cryo-ET experiments of *in vivo Ec* arrays consistently shows the CheA P3 density to be too sparse to determine its position, cryo-ET of *in vitro* lipid-templated *Ec* arrays clearly defines the P3 position. If the P3 domain does engage receptors, this discrepancy may suggest that lipid-templating abrogates this native behavior. Furthermore, it is unclear how the *Td* chemotaxis arrangements affects cooperative behavior in the arrays as linear CheA:CheW strands are only connected through receptor interactions. Investigations into this system may reveal alternative mechanisms for cooperative behavior than what have been reported. Collectively, our results illustrate the importance of investigating transmembrane systems *in situ,* and that examining systems in non-model organisms can lead to new, unexpected advances for understanding the remarkable signaling system of bacterial chemotaxis.

## Acknowledgements

This work is part of the research programme National Roadmap for Large-Scale Research Infrastructure 2017 – 2018 with project number 184.034.014, which is financed in part by the Dutch Research Council (NWO). This work was funded by a grant from the National Institutes of Health: R35GM122535 awarded to BRC, R01AI078958 and R01DE023080 to CLI, and by the European Union under a Marie-Sklodowska-Curie COFUND LEaDing fellowship to ARM. We thank the Netherlands Centre for Electron Nanoscopy (NeCEN) for access to cryo-ET data collection facilities, and NE-CAT at the Advanced Photon Source for access to X-ray crystallography data collection facilities. NE-CAT is supported by NIH/NIGMS awards P30 GM124165 and S10 RR029205.

## Author contributions

A.R.M., D.R.O, K.K., C.L., B.R.C, and A.B. designed research; A.R.M., D.R.O., and K.K. performed research; A.R.M., D.R.O, W.Y., K.K., and A.S. analyzed data; and A.R.M., D.R.O, and A.B. wrote the paper with input from all authors.

## Competing interest

The authors declare no competing interests.

## Supplementary Methods and Materials

### Data availability

The cryo-ET sub-tomogram averages that support these findings are deposited in the Electron Microscopy Data Bank (EMDB) with accession codes EMD-10784, EMD-10785, EMD-10786, EMD-10787, EMD-10788, EMD-10790. The protein x-ray crystallography structure that supports these findings is deposited in the Protein Data Bank (PDB) with accession code 6Y1Y. These depositions will be publicly available with publication.

### Code availability

All custom-made scripts are available to editors and referees upon request. These scripts will be publicly available with publication.

### Bacterial strains, culture conditions, and oligonucleotide primers

*Treponema denticola (Td)* ATCC 35405 (wild-type) was used in this study. The *Td* deletion mutants, Δ2498 and Δ2498Δ2986, were generated in a previous study^2^. Cells were grown in tryptone-yeast extract-gelatin-volatile fatty acids-serum (TYGVS) medium at 37°C in an anaerobic chamber in presence of 85% nitrogen, 10% carbon dioxide, and 5% hydrogen^1^. *Td* mutants were grown with an appropriate antibiotic for selective pressure as needed: erythromycin (50 μg/ml) and gentamycin (20 μg/ml). *Escherichia coli* 5α strain (New England Biolabs, Ipswich, MA) was used for DNA cloning. The *E. coli* strains were cultivated in lysogeny broth (LB) supplemented with appropriate concentrations of antibiotics. The oligonucleotide primers for PCR amplifications used in this study are listed in Table S3. These primers were synthesized by IDT (Integrated DNA Technologies, Coralville, IA).

### Construction of a CheR truncated mutant (ΔCheR_like_)

*TDE1492::ermB* (Fig. S15) was constructed to replace the CheR-like domain (8781-1,308 nt) in *TDE1492 with* a previously documented erythromycin B resistant cassette (*ermB*)^3^. The *TDE1492::ermB* vector was constructed by two-step PCR and DNA cloning. To construct this vector, the 5’ end of *TDE1492* region and the downstream flanking region were PCR amplified with primers P_1_/P_2_ and P_3_/P_4_, respectively, and then fused together with primers P_1_/P_4_, generating Fragment 1. The Fragment 1 was cloned into the pMD19 T-vector (Takara Bio USA, Inc, Mountain View, CA). The *ermB* cassette was PCR amplified with primers P_5_/P_6_, generating Fragment 2. The Fragment 2 was cloned into the pGEM-T easy vector (Promega, Madison, WI). The Fragment 1 and 2 were digested using *NotI* and ligated, generating the *TDE1492::ermB* plasmid. The primers used here are listed in Table S3. To delete *TDE1492,* the plasmid of *TDE1492::ermB* was transformed into *Td* wild-type competent cells via heat shock, as previously described^4^. Erythromycin-resistance colonies that appeared on the plates were screened by PCR for the presence of *ermB* and absence of *TDE1492 (781-1,308 nt)* gene. The PCR results showed that the *TDE1492 (781-1,308 nt)* gene was replaced by *ermB* cassette as expected (Fig. S15). One positive clone *(ΔTDE1492’)* was selected for further study.

### Bioinformatics software and resources

The datasets used in the bioinformatics analysis were built using data from Microbial Signal Transduction Database v3 (MiST3) accessed February 2020^5^ and the Genome Taxonomy Database v89 (GTDB)^6^. We built custom scripts using TypeScript-3.7.5 and NodeJS-12.13. To make these scripts, we also used packages publicly available at the node package manager repository (npm): we used RegArch-1.0.1^7^ to separate CheW-CheR_like_ from other CheWs, gtdb-local-0.0.12 (https://npmjs.com/package/gtdb-local) to use the GTDB taxonomy, Phylogician-TS-0.10.1-4 (https://npmjs.com/package/phylogician-ts) to visualize and manipulate the phylogenetic trees, BioSeq-TS-0.2.4 (https://npmjs.com/package/bioseq-ts) to handle protein sequences and MiST3-TS-0.7.6 (https://npmjs.com/package/mist3-ts) to access MiST3 API. Multiple sequence alignments were produced using L-INS-I algorithm from the MAFFT package. To reduce redundancy in sequence datasets we used CD-HIT v4.6^8^ with unaligned sequences and Jalview^9^ with aligned sequences. RAxML v8.2.10^10^ was used to perform phylogenetic reconstructions, and low support branches in the phylogenetic trees were collapsed with TreeCollapseCL4^11^. Sequence logos were built using Weblogo 3.7^12^.

### Bioinformatics scripts and pipelines

We collected all information on the proteins classified as CheA (96,434) and CheW (134,165). To process this dataset we built several scripts and pipelines to produce the tables, figures and datasets used in this analysis (Fig. S16). The scripts are found in Supplementary File 1.

### Chemosensory profile of Spirochaetotas

The “spiro-pipeline” selects all genomes from Spirochaetota phylum using gtdb-local package to access GTDB v89, then it filters only the genomes that are also present in MiST3 database. It collects the information on MiST for each genome and appends the complete taxonomy information from GTDB and signal transduction profiles. Finally, the pipeline builds the table with the information in markdown (Dataset 2).

### Chemosensory profile of genomes with at least one CheA-F2

The “chea-pipeline” starts from the raw dataset taken from MiST3 database with information on 96,434 CheA genes. Based on MiST3 classification it selects the genomes with at least one CheA-F2 sequences and fetches information about these genomes. At this step, it also checks with the list generated by the “wr-pipeline” of the genomes containing CheW-CheR_like_. It proceeds to append chemosensory information for each genome and GTDB taxonomy. At this step the pipeline splits into four pathways, where one parses the information to build the Dataset 1 and the other three build FASTA formatted files with sequences from: CheA-F2, CheR-F2, CheW-CheR_like_ and all CheWs belonging to genomes with at least 1 CheA-F2. We noticed that MiST3 currently misclassifies some CheR proteins, so we used RegArch to filter out false positives. We also used RegArch to separate CheW and CheW-CheR_like_ sequences as MiST3 classifies both as CheW (scaffold). The RegArch definitions can be found in the script ‘regArchDefinitions.ts’ of the source code in Supplementary File 1.

In MiST3, 306 genomes contain at least 1 CheA-F2 (Dataset 1) with three exceptions: two of them belong to Acidobacteriota phylum,and one from Planctomycetota, which suggest that the presence of an F2 system in these genomes (outside of the Spirochaetota phylum) is the consequence of lateral gene transfer. Of the 306 genomes with at least one CheA-F2, only the following lack the CheW-CheR_like_ protein: the three non-Spirochaetota genomes mentioned previously, 40 genomes of the Brachyspirae class, and three genomes from the order Borreliales (Fig. S2). Interestingly, the Borreliales genomes do appear to have the CheW-CheR_like_ gene, except there is no gene product associated with them in the MiST3 database.

### Classification of CheW

CheW proteins are not classified in MiST3. In order to make comparisons between the sequences of CheW domains in F2 systems, we must first select only CheW-F2. We first assign to the F2 class all the canonical CheW found in genomes with a single CheA-F2. Contrary to the F2 systems where the canonical CheW is not present in the chemosensory gene clusters, other classes do contain their canonical CheW within the rest of the gene cluster. CheW found within 5 genes from a classified CheA were assigned to the same class as CheA. To perform this classification, we selected the 598 full-length CheW sequences generated by the chea-pipeline. We used CD-HIT to remove 405 redundant sequences (-c 1). Next, we ran the “classify-w” pipeline on the remaining sequences (193). The pipeline reads the identifiers of the sequences and fetches the chemotaxis profile from MiST3 for each genome. It classifies CheWs as F2 classes if there is only 1 CheA of the class F2 in the profile. Next, it fetches the gene neighborhood (5 genes up and downstream) of the remaining CheWs and assigns a matching class if there is a classified CheA within these genes. We also aligned the sequences (193) with the L-INS-I algorithm of the MAFFT package, produced a phylogeny with 1000 rapid bootstrap using RAxML (-f a -m PROTGAMMAIAUTO -N 1000) and collapsed the phylogeny using TreeCollapse4 at 50% bootstrap. We mapped the CheW classification to the CheW tree in Figure S2. We expanded the F2 classification to the 74 sequences within the branch with only CheW-F2 sequences.

### Comparison of the CheW domains in CheA, CheW-CheR_like_ and CheW

We put together the sequences from CheA-F2 and CheW-CheR_like_ (both trimmed by the PFAM model for the CheW domain, already annotated in MiST3) and the full sequence of the CheW-F2 selected in the previous step. We then use L-INS-I algorithm to align the sequences and Jalview to manually inspect and eliminate identical redundant sequences. Finally, we trimmed the whole alignment based on the boundaries of CheA-F2 and CheW-CheR_like_ and eliminated one incomplete sequence: GCF_000413015.1-HMPREF1221_RS07250. The final alignment had a total of 206 sequences: 73 CheA, 59 CheW-CheR_like_ and 74 CheW. We separated the alignments into individual files and built sequence logos to summarize the amino-acid diversity in each position for each group (Fig. S3B).

### Comparison of CheR domains

First, we added together the trimmed part matching the CheR domain of the CheW-CheR_like_ sequences and the 292 sequences of the CheR protein. We then aligned the 550 sequences using L-INS-I. We used Jalview to inspect the alignment and remove identical redundant sequences. The final alignment had the CheR domain of 83 CheR and 83 CheW-CheR_like_ proteins. We separated the alignments into individual files and generated independent sequence logos (Fig. S3A).

### Analyses of CheA P3 domains

The “p3-pipeline” pipeline processes the data for the analysis of the length of P3 domains of all CheA homologs in the MiST3 database. It reads the information for all 96,434 CheAs in the MiST3 database, trims the sequence matching the Pfam mode H-kinase_dim and builds FASTA formatted datasets for each chemotaxis class. For each dataset, we used CD-HIT with 75% identity cutoff and aligned them using the L-INS-I algorithm from MAFFT. Using Jalview we manually inspect and edited the alignment to remove divergent sequences that opens major gaps in the alignment. We removed 6 F1 sequences, 37 F7 sequences, 20 F8 sequences and 5 F9 sequences. Then we merged the alignment using mafft-profile with each dataset as a seed alignment in a single shot (Fig. S13A). We selected the non-conserved central region of the alignment and measured the number of amino-acids in each sequence (Fig. S13B).

### Cryo-ET and sub-tomogram averaging of *T. denticola* chemotaxis arrays

Cells were concentrated by centrifugation, and a 1/10 dilution of protein A-treated 10-nm colloidal gold solution (Cell Microscopy Core, Utrecht University, Utrecht, The Netherlands) was added to the cells and mixed by pipeting. 3 μL aliquots of the cell suspension were applied to glow-discharged R2/2 200 mesh copper Quanti-foil grids (Quantifoil Micro Tools, GmbH), the sample was pre-blotted for 30 seconds, and then blotted for 2 seconds. Grids were pre-blotted and blotted at 20 °C and at 95% humidity. The grids were plunge-frozen in liquid ethane using an automated Leica EM GP system (Leica Microsystems).

Data collection was achieved on a Titan Krios transmission electron microscope (Thermo Fisher Scientific) operating at 300 kV. Images for three strains (WT, Δ2498, Δ2498Δ2986) were recorded with a Gatan K2 Summit direct electron detector with a GIF Quantum energy filter (Gatan) operating with a slit width of 20 eV. Images were taken at a magnification of 42,000×, which corresponds to a pixel size of 3.513 Å. Tilt series were collected using SerialEM with a modified bidirectional tilt scheme (−20° to 60°, followed by −22° to −60°) with a 2° increment. Images for the ΔCheR_like_ strain were recorded with a Gatan K3 Summit direct electron detector equipped with a GIF Quantum energy filter (Gatan) operating with a slit width of 20 eV. Images were taken at a magnification of 26,000X, which corresponds to a pixel size of 3.27 Å. Tilt series were collected using SerialEM with a bidirectional dose-symmetric tilt scheme (−60° to 60°, starting from 0°) with a 2° increment. For all strains, the defocus was set to – 6μm and the cumulative exposure per cell was 100 e-/A^2^.

Bead tracking-based tilt series alignment and drift correcting were done using IMOD^13^. CTFplotter was used for contrast transfer function determination and correction^14^. Tomograms were reconstructed using simultaneous iterative reconstruction with iteration number set to 6. Dynamo was used for manual particle picking and sub-tomogram averaging^15,16^.

### Purification of CheA, CheA P3, CheW-CheR_like_, and TDE2496 proteins of *T. denticola*

DNA segments encoding the CheA P3 domain, CheW-CheR_like_ protein, and TDE2496 in *T. denticola* were PCR amplified from *Td* genomic DNA using a forward oligonucleotide encoding an NdeI restriction site and a reverse primer encoding an EcoRI restriction site. The CheA protein was amplified using a forward oligonucleotide encoding an NdeI restriction site and a reverse primer encoding an BamHI restriction site. The PCR products were treated with the appropriate restriction enzymes, purified, and ligated into a pet28a plasmid with a poly-Histidine tag and kanamycin resistance marker. The plasmids were transformed into *Escherichia coli* BL21-DE3 cells and 4-8 L of cell culture were grown at 37°C until an O.D. of 0.6 was reached. The flasks were cooled to 21°C and 1 mM of IPTG was added to the culture. The cells were harvested after 16 hours of growth. The cells were lysed via sonication in lysis buffer (50 mM Tris pH 7.5, 150 mM NaCl, 5 mM Imidazole) while cooled on ice. The lysate was centrifuged at 20,000 X G for 1 hour at 4°C. The lysate was then run over a gravity-flow purification column containing 3 ml of Nickel-NTA resin. The resin was washed with 10 ml wash buffer (50 mM Tris pH 7.5, 150 mM NaCl, 20 mM Imidazole) and the protein was eluted with 10 ml elution buffer (50 mM Tris pH 7.5, 150 mM NaCl, 200 mM Imidazole) and collected in 1 ml fractions. The fractions were assessed for protein concentration via Bradford reagent and the fractions containing protein were run on a size-exclusion s75 and s200 column systems that monitored absorbance at 280 nm and collected 6 ml fractions. Fractions that contain CheA were concentrated to ∼20 mg/ml via centrifugation in a protein concentrator containing a regenerated cellulose filter with a 50 kDa molecular-weight cut-off (MWCO). Fractions that contain CheA P3, CheW-CheR_like_ and TDE2496 were concentrated with a 10 kDa MWCO filter to 32 mg/ml, 11 mg/ml and 7 mg/ml, respectively. The protein solutions were aliquoted, flash frozen in liquid nitrogen, and stored at −80°C. For CheW-CheR_like_, CheA, and CheA P3 domain, the purifications were prepared at ambient temperatures. For TDE2496, the purification was prepared at 4°C.

### Radioisotope assays

23 μl samples containing 2 μM CheA alone, or 2 μM CheA and 2 μM CheW-CheR_like_, or 2 μM CheA and 2 μM CheW-CheR_like_ with TDE2496 (2 μM or 12 μM), were incubated in 50 mM MOPS pH 7.5, 150 mM KCl, 10 mM MgCl_2_ for 30 minutes at ambient temperatures. Phosphorylation of CheA was initiated by the addition of 2 μl of a solution containing 1 mM ATP mixed and 2-10 μl radiolabeled γ-P32 ATP (3000 Ci/mmol, 10 μCi/μL; Perkin Elmer) and quenched with 25 μl of 3X LDS buffer containing 100 mM EDTA pH 8.0 after 1-12 minutes. The samples were run on a native Tris-glycine gel for 2 hours at 120 volts. The gels were dried, placed in a radiocassette for 24 hours, and then imaged with a Typhoon phosphor-imager (GE Healthcare). The intensity of the radio-labeled protein bands were quantified using ImageJ.

### Quantification of cell curvature

The cell curvature of *Td* whole cells and *V. cholera* minicells was quantified by analyzing images of cross-sections of the cells where top views of chemotaxis arrays are present. For *Td* cells, the inner membrane curvature and CheA:CheW base-plate curvature was quantified. For *Vc* minicells, the inner membrane curvature was quantified. These images were pre-processed with Fiji by placing points along the desired area with a distance of 10 nm between each point. The curvature of the inner membrane was measured with a pre-built plugin for python, called Sabl_mpl, written by Jewett, A. from the Jensen lab (Pasadena, CA)^23^. The “measure 3-point curvature” function was used to select three adjacent points along the inner membrane of the cell and calculate the radius of these points. The radius of the three selected points allowed for the calculation of the local curvature by dividing 1 with the measured radius (1/R = c). This was repeated for all points with a “sliding window” approach, where the second point of the initial three points would become the first point, until the desired area was covered.

### Quantification of array alignment to the cell axis

The angle between the strands of CheA:CheW rings and the *Td* cell axis was quantified using Image J software. 2D images from reconstructions that clearly locate the orientation of the strands and cell axis were chosen for analysis. First, a straight line was drawn from the cell pole down the axis of the cells. Then, a second line was drawn through one of the strands in the array and the angle between the two intersecting lines was quantified. In some cases, the angle was too small (<3°) to be accurately determine so the angle was annotated as 0°.

### Residue conservation and molecular modeling of *Td* CheW, CheA P5, and the CheW domain of CheW-CheR_like_

The protein sequences of the two *Td* CheW domains and *Td* P5 were aligned (Clustal Omega), conservation was calculated based on the alignment (JalView^9^), and the highest variable regions were selected based on conservation (10+ adjacent residues with conservation scores lower than 8). Homology models of *Td* CheA P5, CheW and the CheW domain of CheW-CheR_like_ were generated via the Swiss-Model server using complete residue sequences of each protein as a target^24^. The CheW protein from *Thermoanaerobacter tengcongensis* (*Tt*) (PDB ID: 2QDL) had the highest percent identity to the CheW proteins (37% and 31%) and was therefore used as the structural template. The resulting homology models for *Td* CheW and the CheW domain of CheW-CheR_like_ had a QMEAN of – 1.3 and −1.2, respectively. The P5 structure from *Escherichia coli* produced the best homology model for *Td* CheA P5, with QMEAN −0.98 (PDB ID: 6S1K). The homology models were then aligned to the CheW protein and P5 domain in a crystal structure containing *Thermotoga maritima* CheW and CheA P4P5 in complex using PyMol (PDB ID: 3UR1).

### Crystallization and structural determination of the P3 domain of *T. denticola* CheA

The isolated P3 domain was concentrated to 32 mg/ml and crystallized via hanging drop in 0.1 M Imidazole pH 7.0, 25% PEG 400 using a 1:1 ratio of protein solution to crystallization solution with a final volume of 3 μl. Crystals were apparent within eight hours but increased in size over three days. Crystals were manually picked up in loops, flash cooled and shipped in liquid nitrogen to a beamline (APS, line NE-CAT 24-ID-C, Dectris Pilatus 6M-F Pixel Array detector). The crystals diffracted to ∼1.3 Å with a C2 symmetry and data was cut-off at 1.5 Å. The diffraction data was scaled and integrated using XDS^17^, and phased by molecular replacement with Phaser MR using *ab initio* search models generated through the QUARK server and then ran through the AMPLE pipeline on the CCP4 web server^18–20^. Model improvement was done by several rounds of manual model improvement in COOT followed by automated refinement using Phenix Refine software^21,22^.

---

**Fig. S1.**
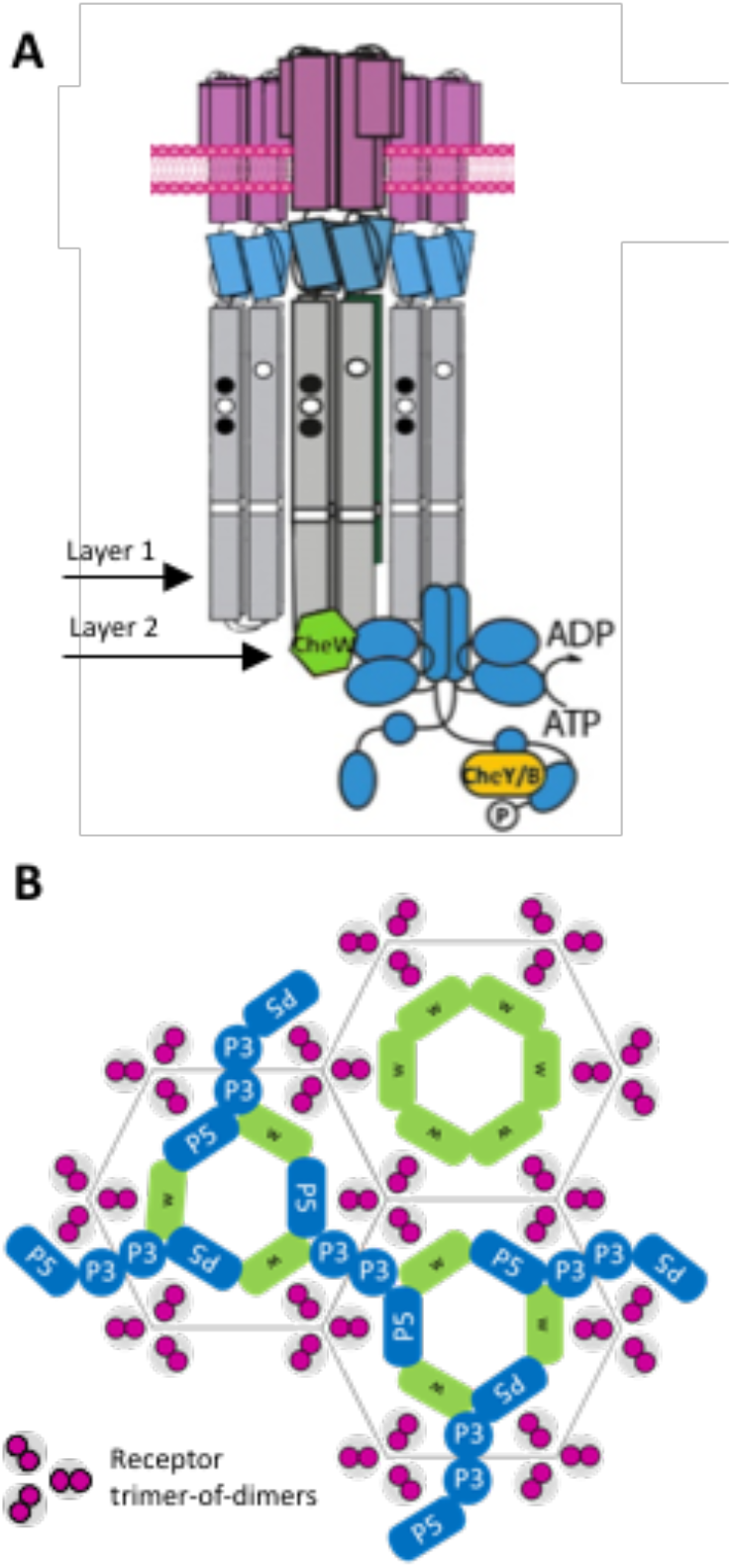
The arrangement of the canonical chemoreceptor array, exemplified in *Escherichia coli* (*Ec*). (A) Transmembrane chemoreceptors form dimers that function as a trimer-of-dimers module to modulate the activity of the histidine kinase CheA (blue) via a coupling protein CheW (green). In cryo-ET experiments, crosssections at Layer 1 reveals the arrangement of chemoreceptors and Layer 2 reveals the position of the kinase. (B) In all organisms investigated previously, CheA is integrated into the array with hexagonal (pseudo-P6) symmetry.

**Fig. S2.**
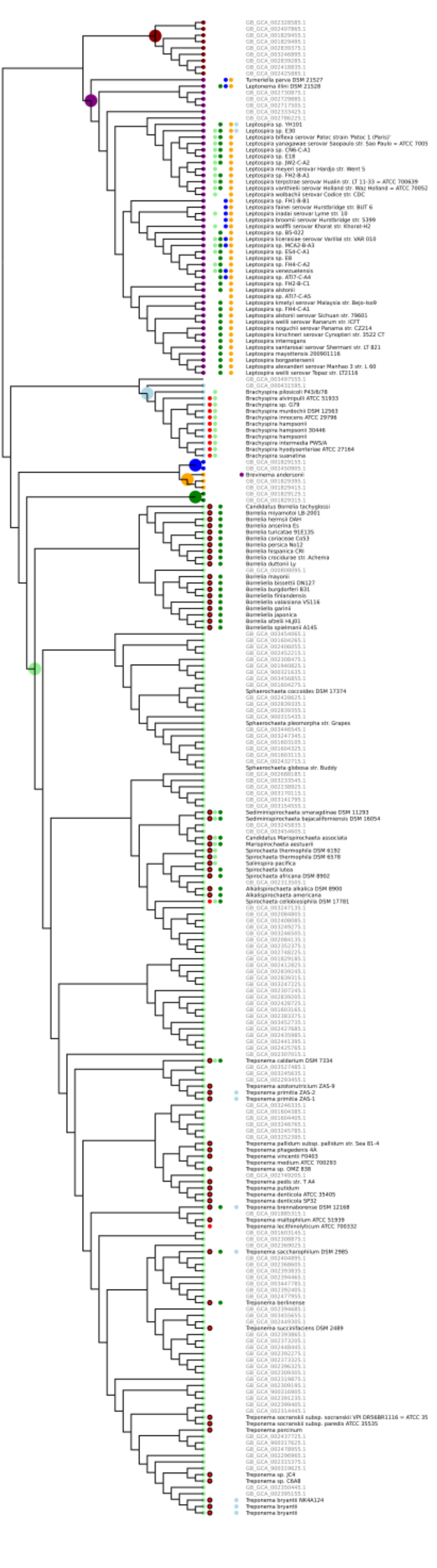
The chemosensory class profile of Spirochaetota. Sequences belonging to the four major classes of Spirochaetota are marked: UBA4802 (dark red), Leptospira (purple), Brachyspirae (cyan), UBA6919 (blue), Brevinematia (yellow), GWE2-31-10 (green) and Spirochaetia (light green). Nodes are marked for presence of classes F1 (orange), F2 (red), F5 (blue), F7 (light green), F8 (green), ACF (light blue), and Tfp (purple). Genomes not present in MiST3 database are marked as grey. F2 systems with CheW-CheR_like_ are marked with a black outline around the red circle.

**Fig. S3.**
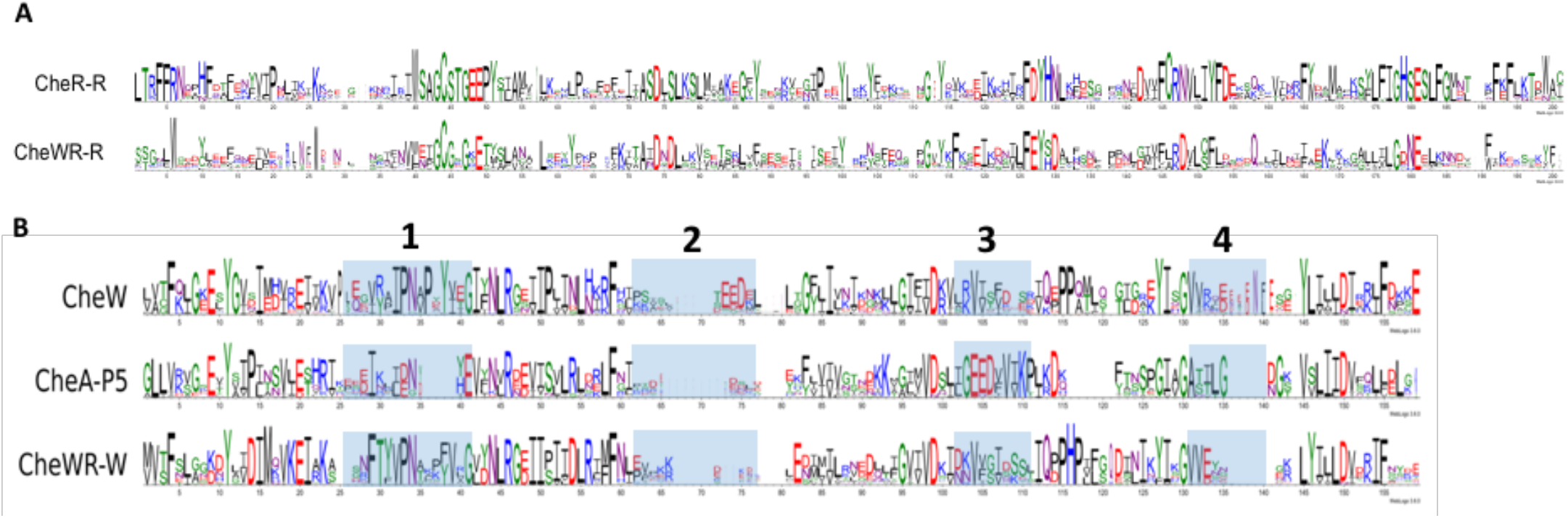
(A) Sequence logo of representative sequences of the two groups of CheR-containing proteins in the F2 system. (B) Sequence logo of representative sequences of the three groups of CheW-containing proteins in the F2 system. Blue boxes denote the location of variable regions identified in the *Td* CheW and P5 homologs. In these locations, all regions possess unique conserved residues with the exception of Region 2, which is not at the CheA:CheW ring interfaces.

**Fig. S4.**
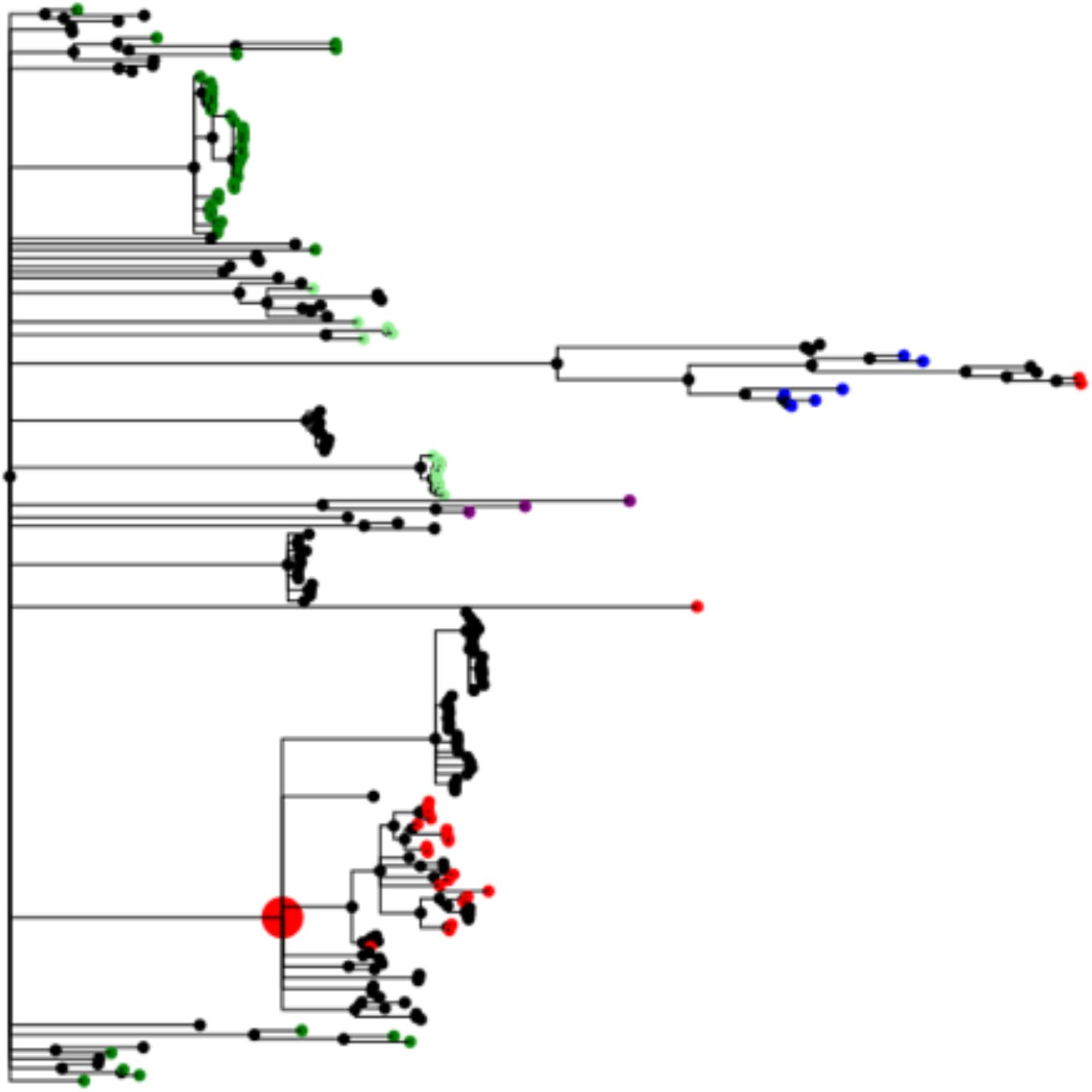
Phylogenetic tree of CheW sequences in genomes with at least one CheA-F2 suggests a last common ancestor of CheW-F2 sequences. We mapped the sequences of CheW from the classes F2 (red), F5(purple), F7 (light green), F8 (green) and ACF (blue). The larger red internal node marks a candidate of the last common ancestor of CheW-F2.

**Fig. S5.**
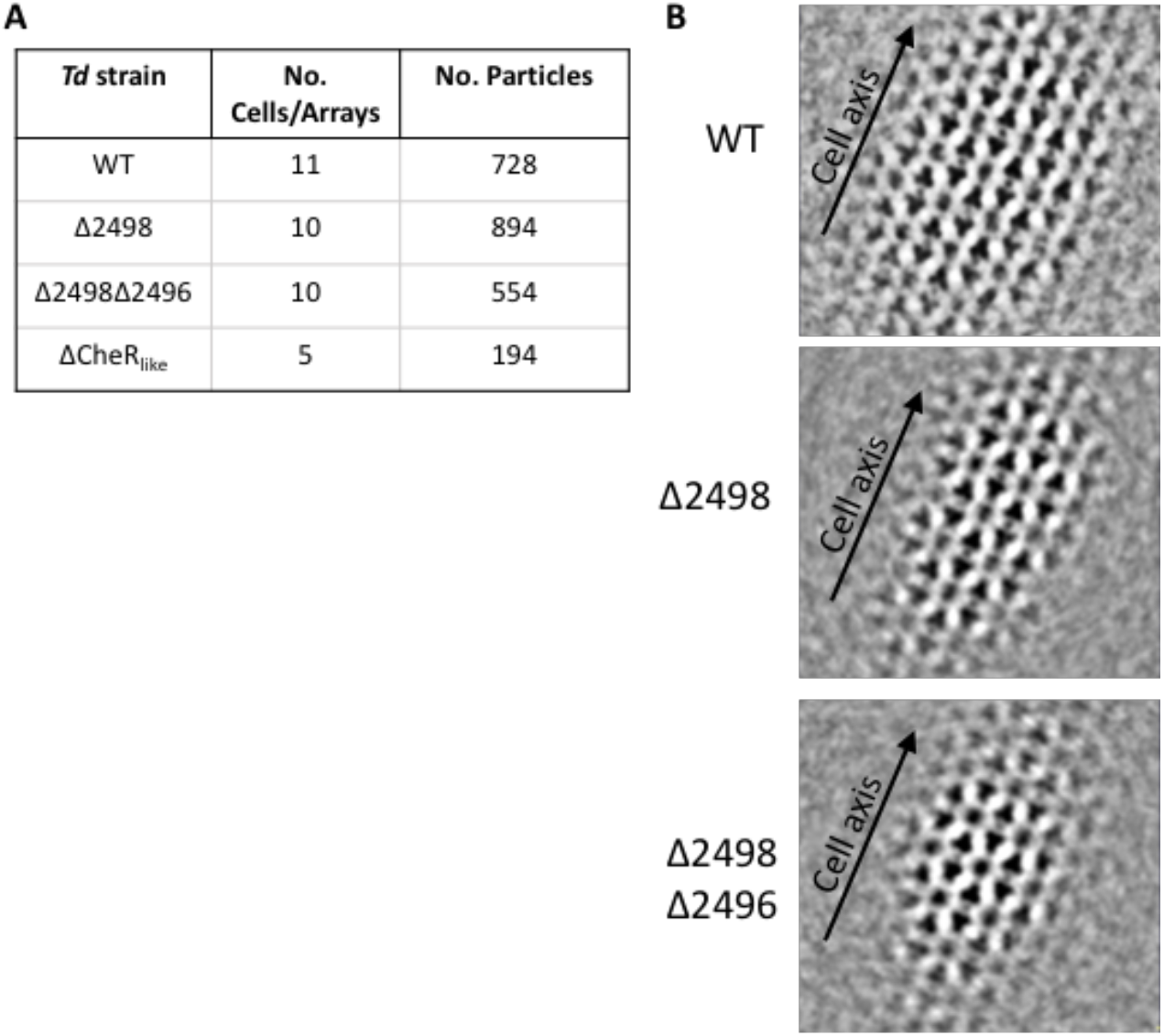
(A) The number of arrays and particles selected to generate subtomogram averages for each *Td* strain. (B) Sub-tomogram averages reveal the orientation of the cell axis with respect to the orientation of the chemotaxis arrays. Intriguingly, the arrays have a preferred orientation in the cells and this orientation is conserved in WT and deletion mutants.

**Fig. S6.**
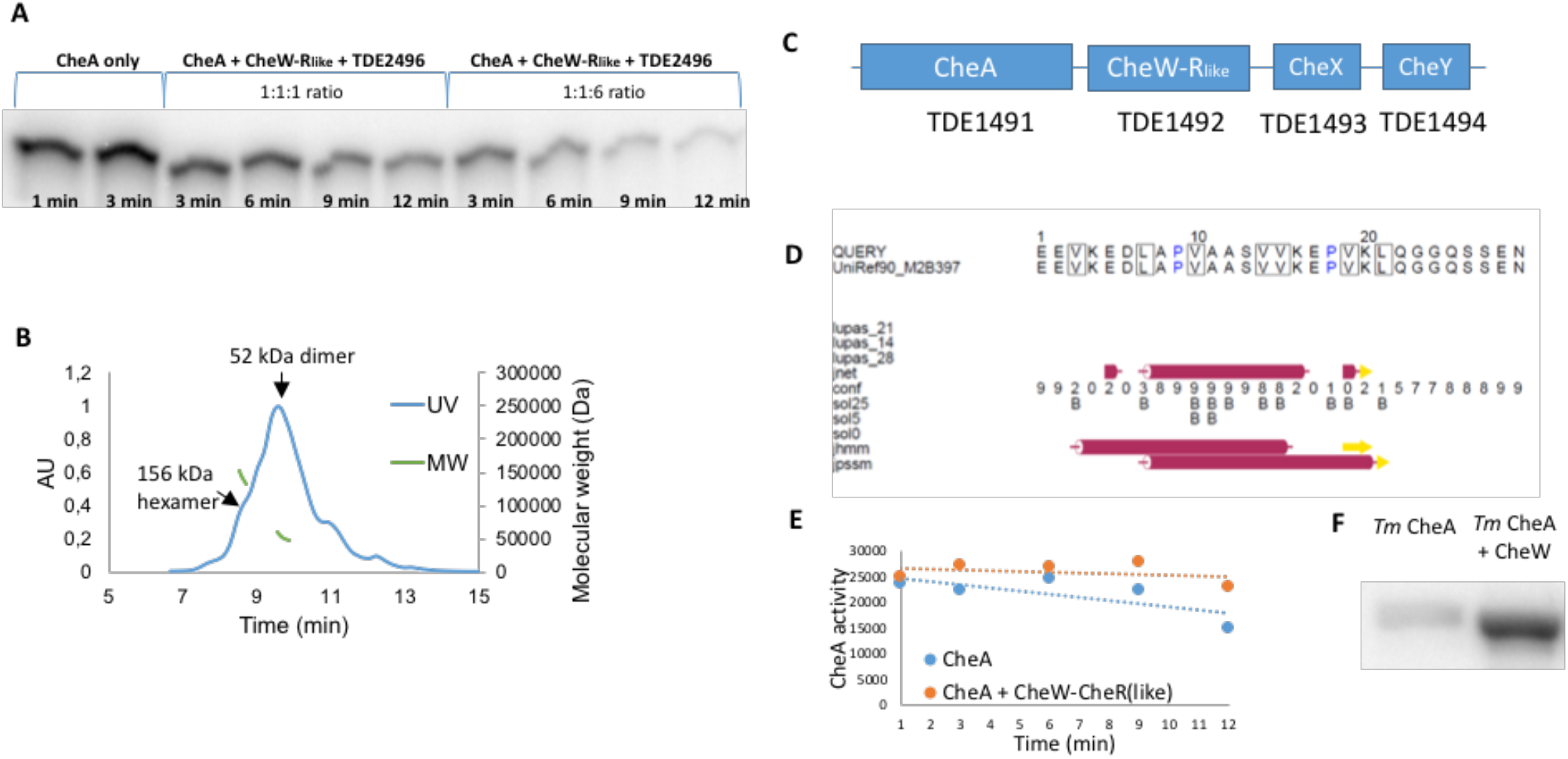
(A) The soluble receptor TDE2496 deactivates *Td* CheA in a concentration-dependent manner. (B) MALS experiments demonstrate that TDE2496 forms the expected dimer (52 kDa) and hexamer (156 kDa) oligomeric states. (C) The CheW-CheR_like_ protein is conserved on the same operon as the only CheA, CheX, and CheY proteins in *Td.* (D) The CheW-CheR_like_ linker is predicted to forma single alpha helix flanked by unordered regions (Jpred). (E) Radioisotope assays that monitor CheA autophosphorylation over time in the presence and absence of CheW-CheR_like_ demonstrate a difference in CheA activity. (F) In the *T. maritima* system, the presence of CheW also activates CheA kinase activity. Both samples are at time point 3 minutes.

**Fig. S7.**
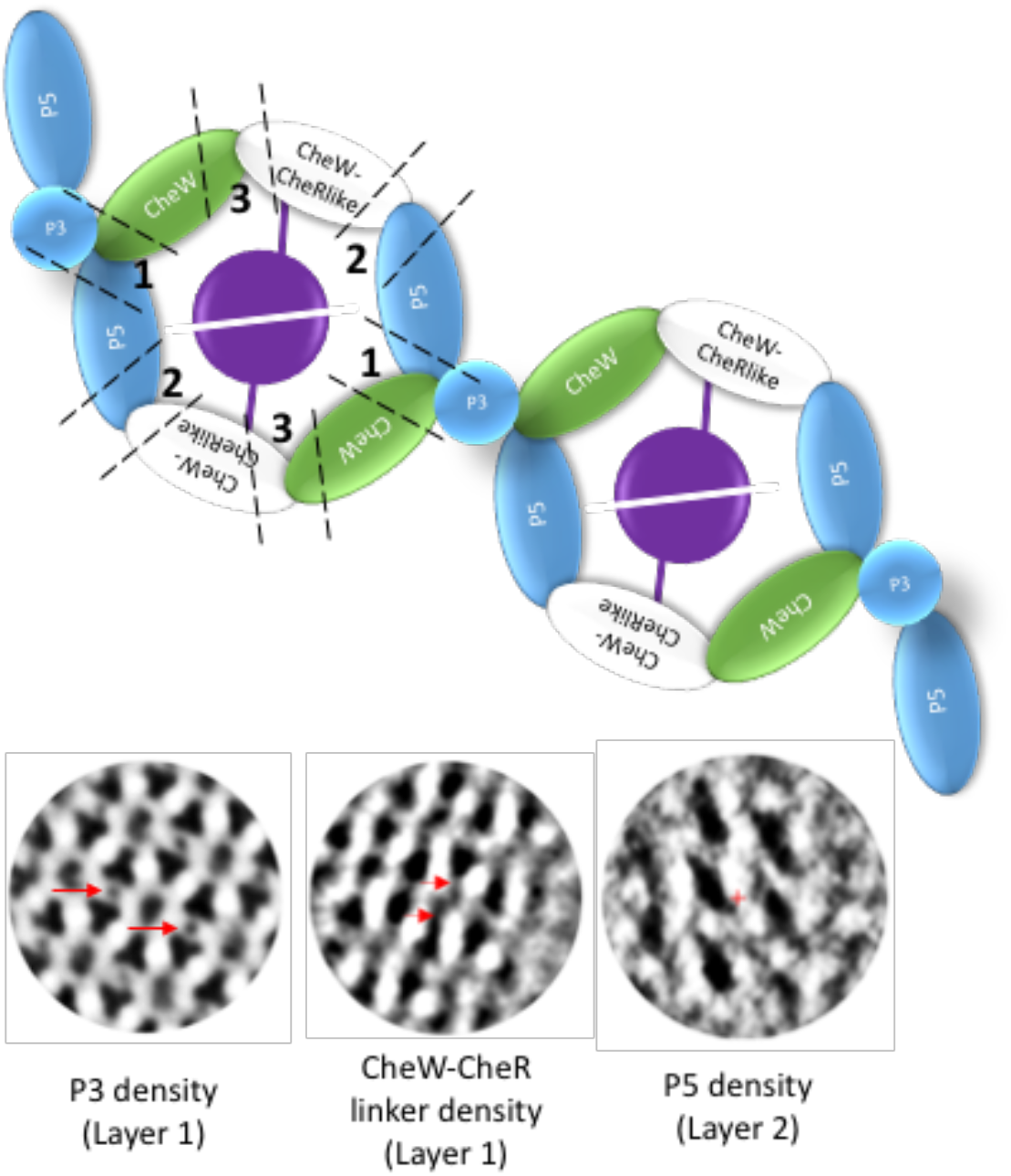
The arrangement of chemotaxis CheA:CheW rings in *Td.* Top: The CheA:CheW rings consists of three domains: CheA P5 (blue), a classical CheW protein (green), and the CheW domain of CheW-CheR_like_ (white). Three unique interfaces are produced and the CheR_like_ domain (purple) extends to the center of the rings to interact with one another. Bottom: The location of the CheW-CheR_like_ protein in the rings is evident by examining the location of the CheA domains with respect to the CheW-CheR_like_ linker.

**Fig. S8.**
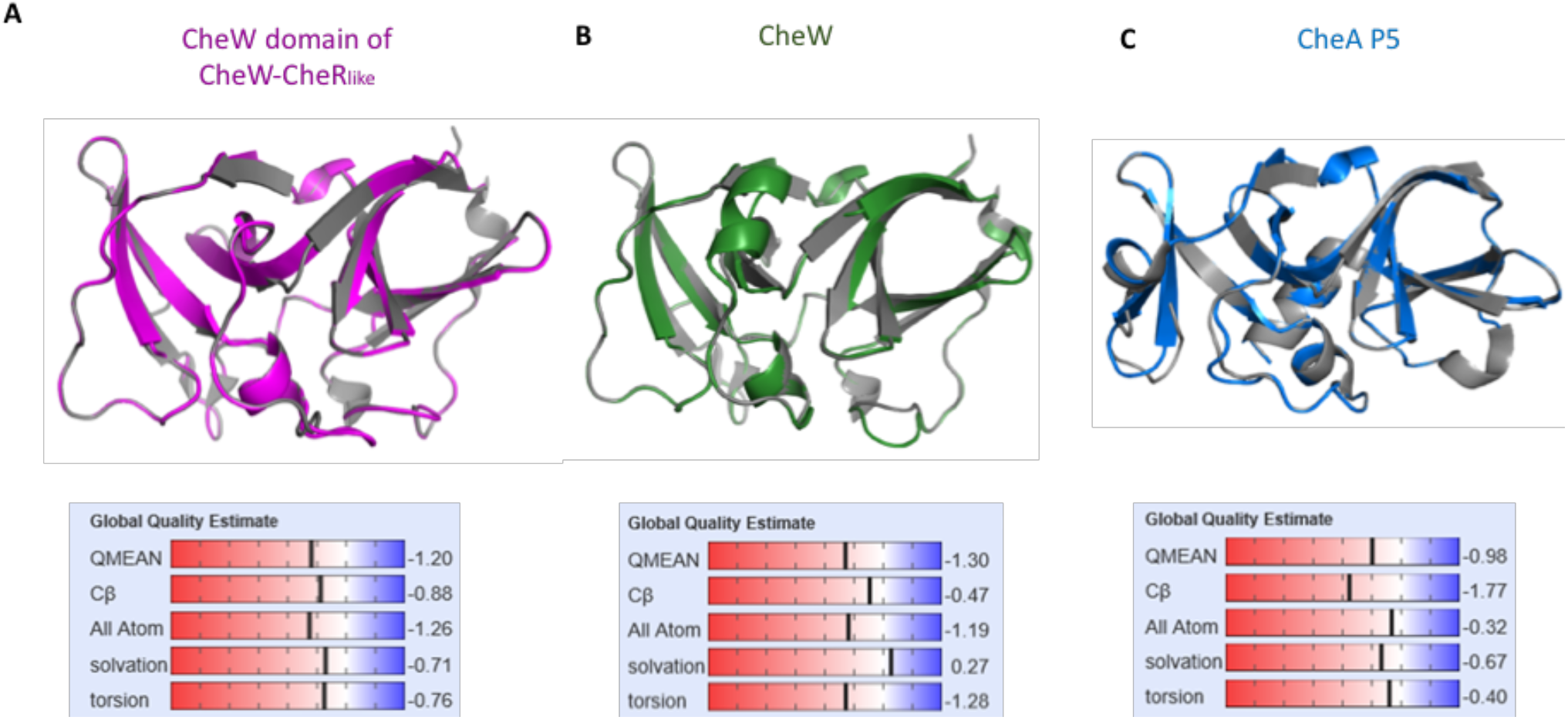
Homology models of *Td* CheW domains and CheA P5. (A) A homology model of the CheW domain of CheW-CheR_like_ using *Thermoanaerobacter tengcongensis* (*Tt)* CheW (PDB ID: 2QDL) as the template. (B) A homology model of the classical *Td* CheW using *Tt* CheW (PDB ID: 2QDL) as the template. (C) A homology model of CheA P5 using *E. coli* CheA P5 (PDB ID: 6S1K) as the template.

**Fig. S9.**
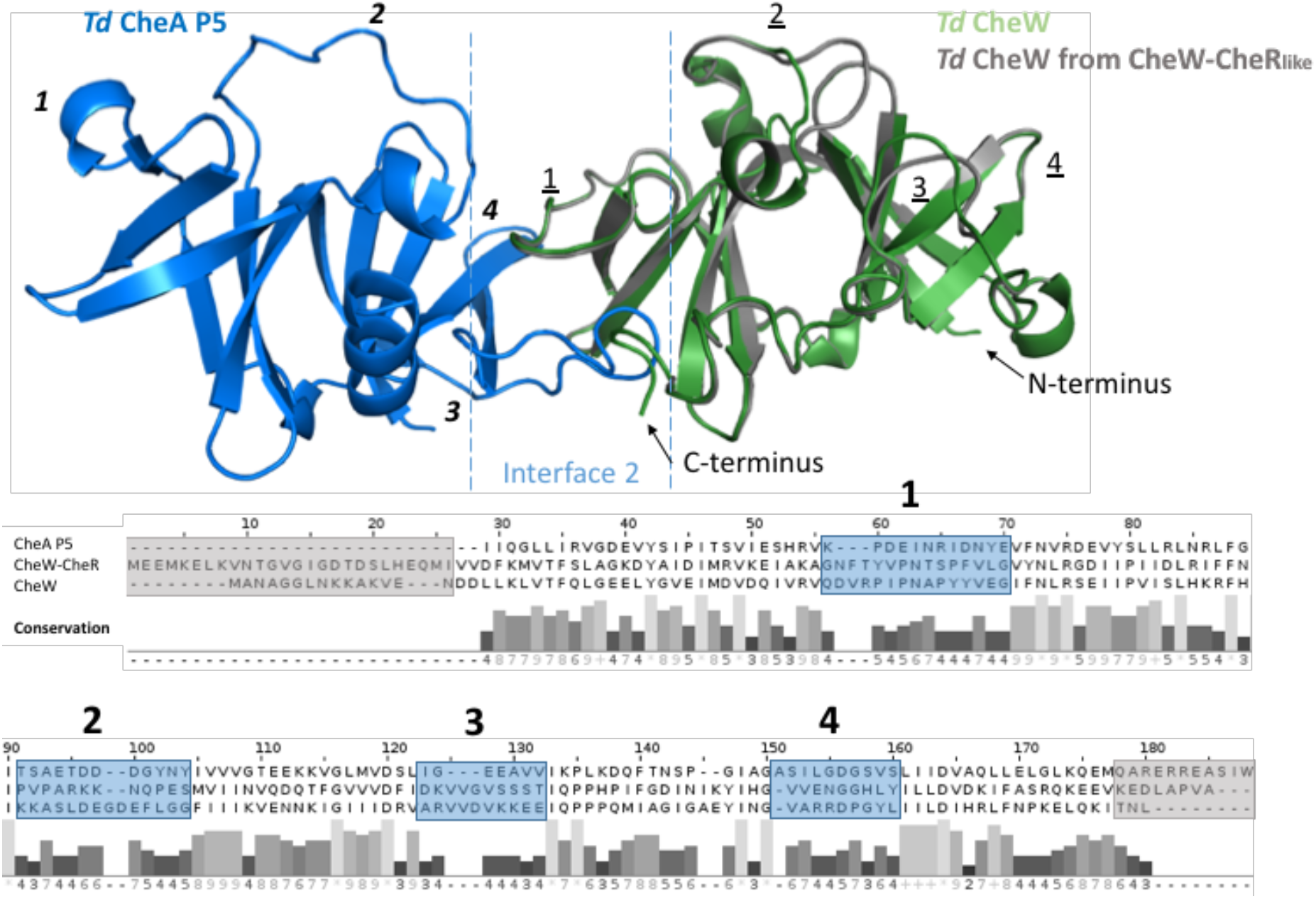
Homology models of *Td* CheA P5, CheW, and the CheW domain of CheW-CheR_like_ mapped onto a previously determined crystal structure of *Thermotoga maritima* CheA P5 and CheW (PDB ID: 3UR1). Variable regions between the three domains (blue boxes) were determined by sequence alignments of the three domains followed by conservation analysis. These variable regions are located at the rings interface regions. The CheW domains also have variable N-terminal and C-terminal regions that are not complimented in P5 (grey boxes). Variable regions for P5 and the CheW domains are denoted on the homology models by italicized numbers and underlined numbers, respectively.

**Fig. S10.**
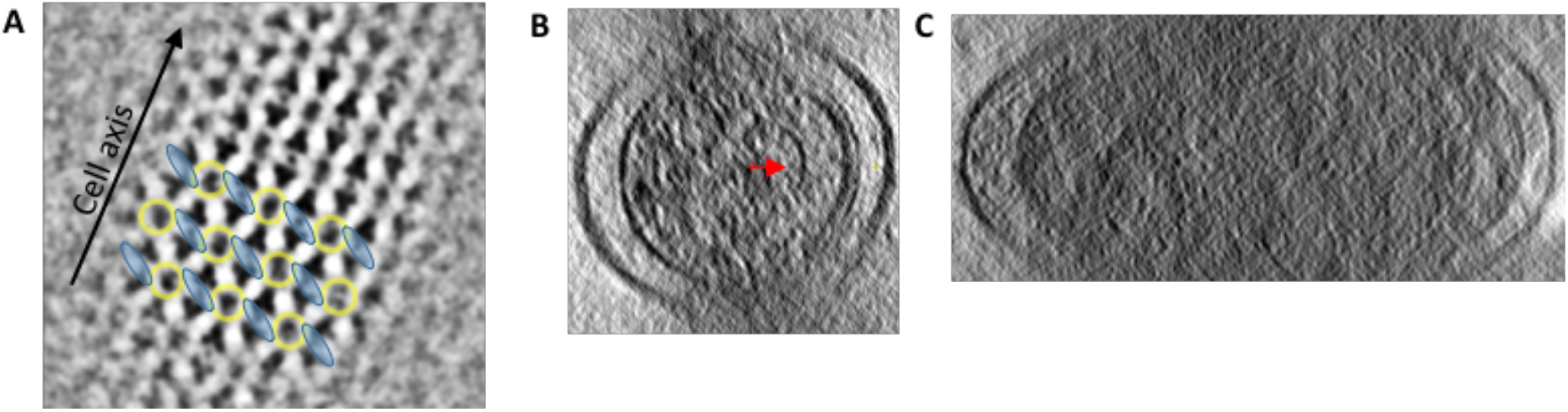
(A) Sub-tomogram averaging of *Td* WT and mutant strains reveal that linked CheA:CheW rings (yellow) run perpendicular to the cell axis via a strict linear orientation of CheA (blue). Averages from WT *Td* are illustrated here but apply to all strains. (B) The curvature of the inner membrane of *Td* at chemotaxis arrays in the reconstructions is 35.8 +/− 6.6 /μm (270 Å radius). The curvature of the CheA:CheW baseplate (red arrow) is 65.6 +/− 19 /μm (152 Å radius). (C) *V. cholerae* minicells have an inner membrane curvature of 9.15 +/− 4.5 /μm (radius 1092 Å).

**Fig. S11.**
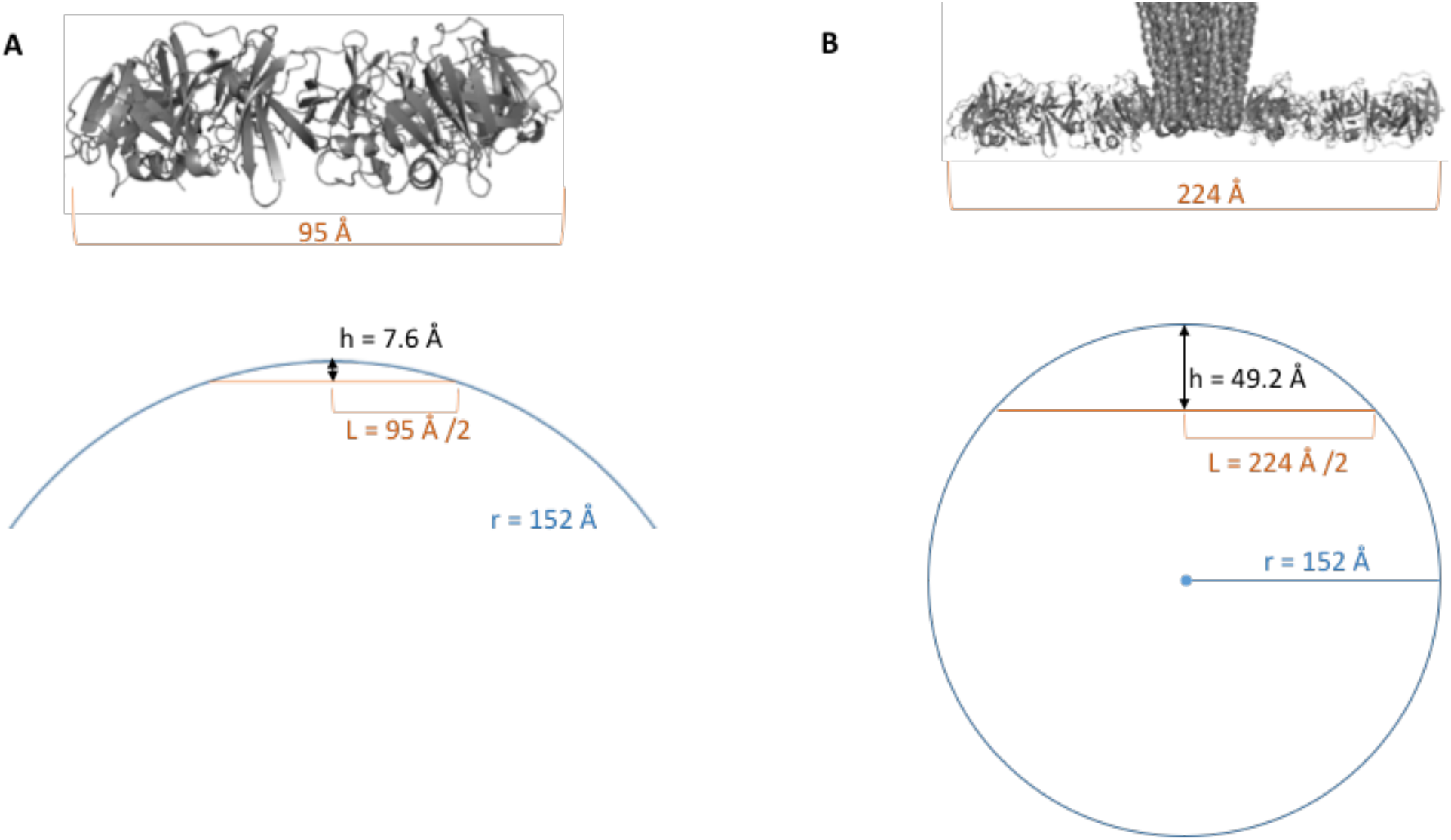
Modeling of CheA:CheW rings to the curvature of the *Td* baseplate. (A) For single CheA:CheW rings to follow the baseplate curvature, it must buckle by an average of 7.6 Å toward the membrane. (B) In order for two linked CheA:CheW rings to run perpendicular to the cell axis, the center of the rings (P3) must buckle by 49.2 Å toward the cell membrane.

**Fig. S12.**
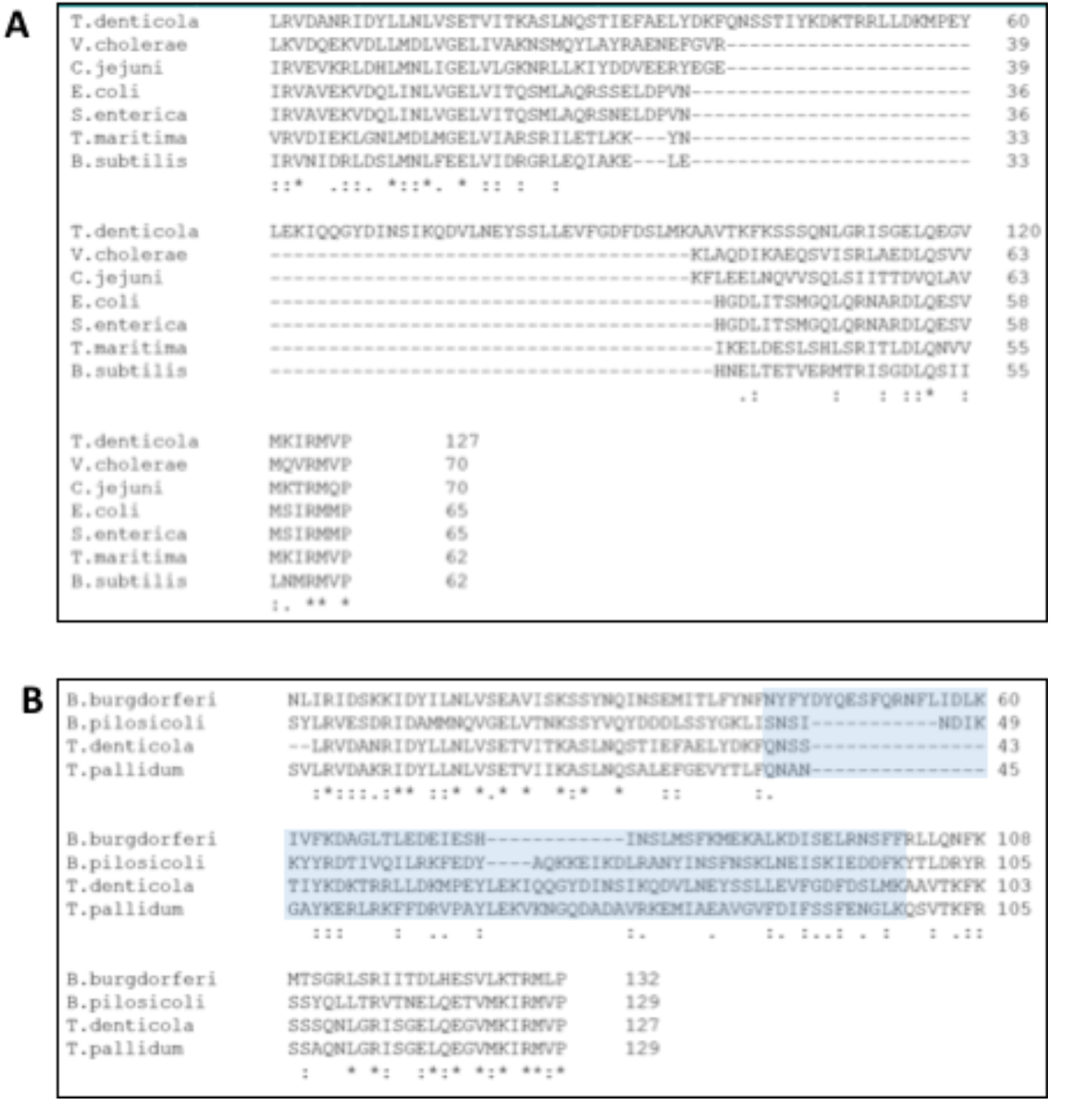
Multiple sequence alignments of the CheA P3 domain from several bacteria demonstrates the presence of additional P3 residues in *Td* and other Spirochetes. (A) CheA P3 alignments of *Td* and P3 from other bacteria with previously characterized chemotaxis proteins. *Td* possesses ∼50 residues that are not found in the other homologs and are located in between the traditional dimerization helices. (B) *Td* CheA P3 aligned with other Spirochete P3. The additional residues identified in alignment **A** are highlighted in blue. Figures were made using Clustal Omega.

**Fig. S13.**
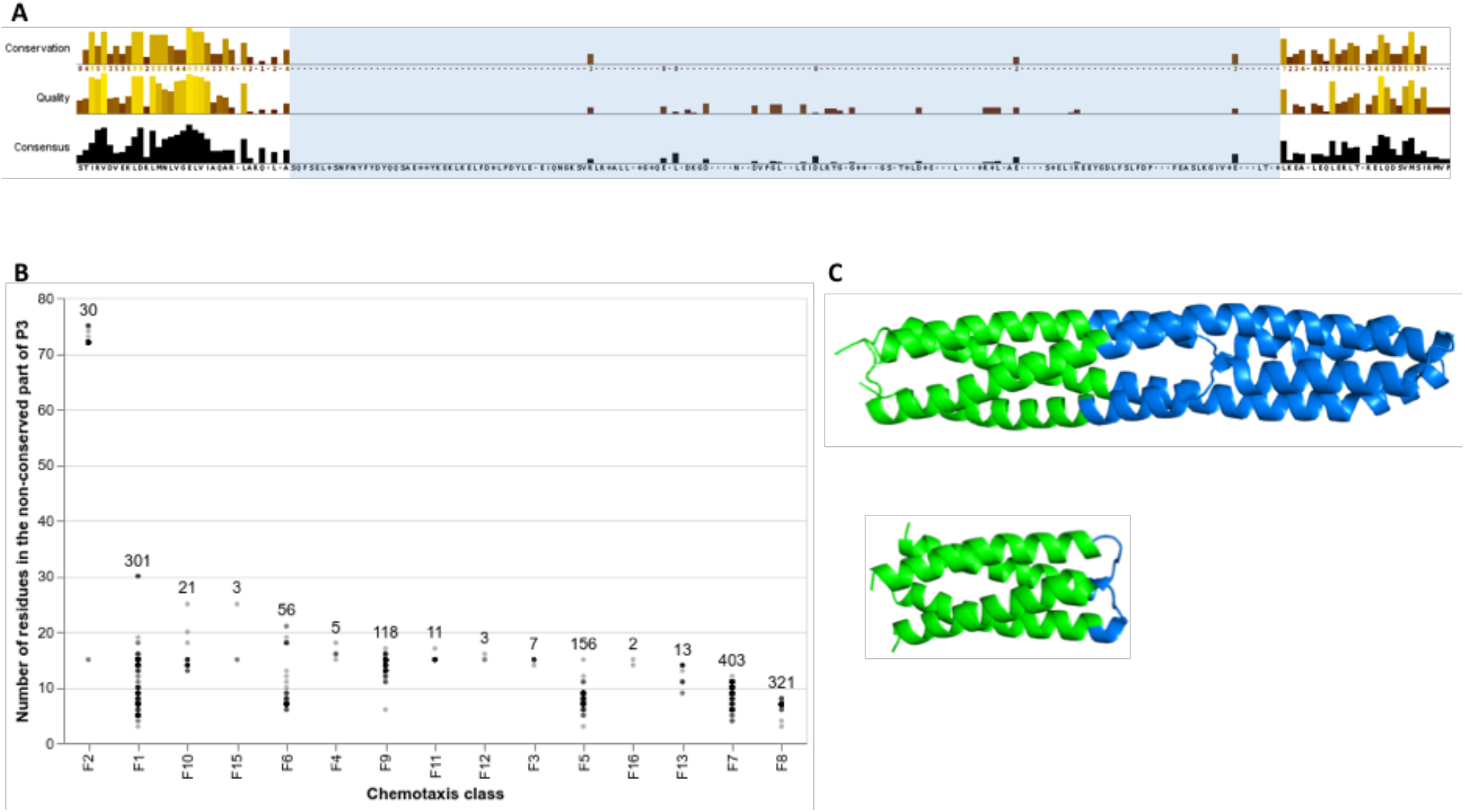
Analysis of non-redundant CheA P3 domains with a 75% sequence identity cut-off (1450 sequences). (A) Conservation scores of the P3 alignment with the 1450 sequences. The traditional P3 helices are largely conserved, but regions located between the helices are non-conserved. (B) Analyses of CheA from different chemotaxis classes reveals that CheA F2 homologs possess the most residues in the non-conserved region of P3. (C) The non-conserved regions are llustrated with the two known CheA P3 structures. Top: *Td* P3 possesses 71 residues that align to the nonconserved region (PDB ID: 6Y1Y). Bottom: *Tm* P3 possesses seven residues that align to the non-conserved region (PDB ID: 1B3Q).

**Fig. S14.**
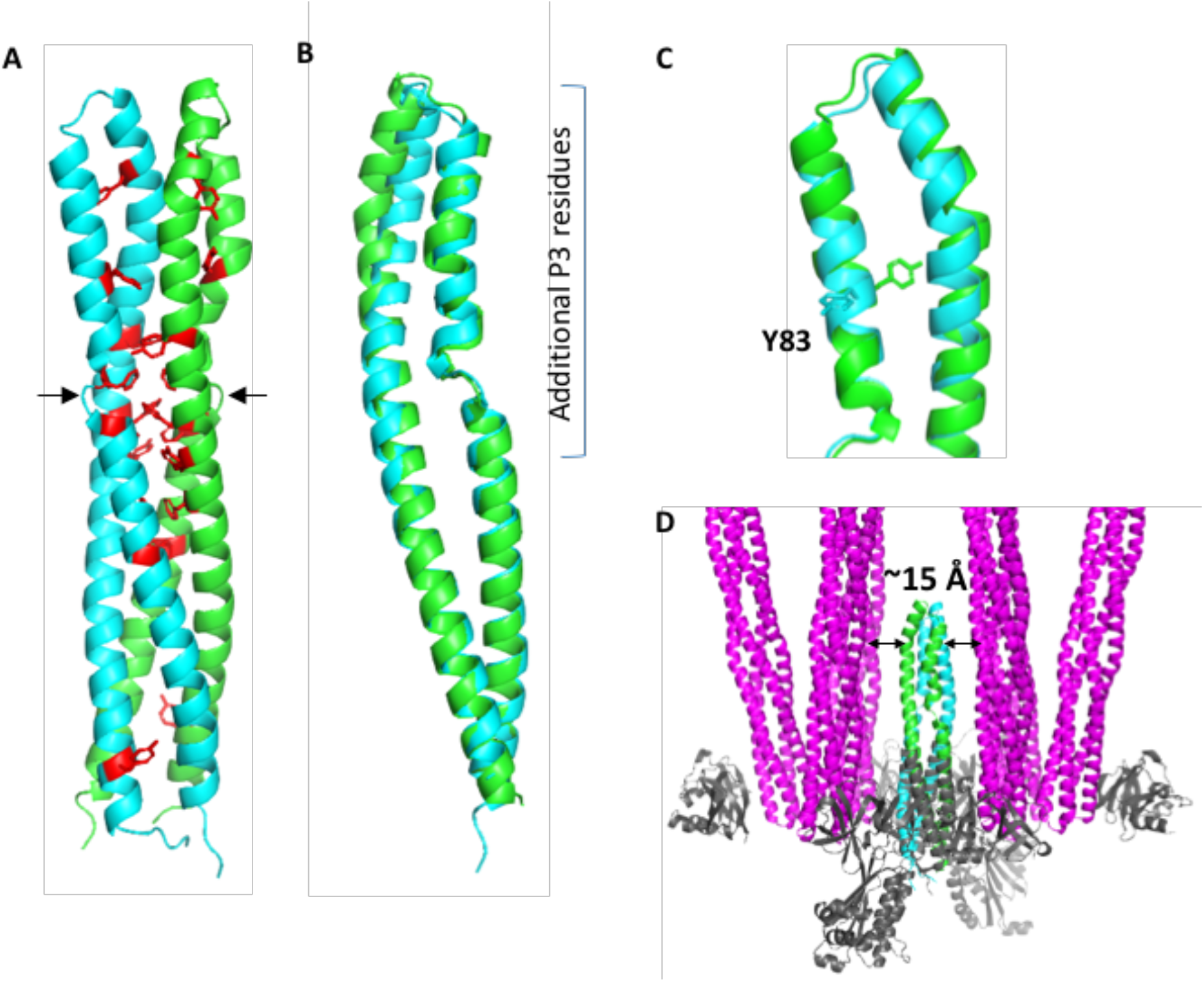
The P3 domain of *Td* CheA. (A) The CheA P3 dimer contains a cluster of Phe and Tyr residues near the breakages in the helices (black arrows). All Phe and Tyr residues are highlighted in red. (B) The crystal structure of CheA P3 demonstrates asymmetry in the subunits. (C) Repositioning of Y82 in the subunits induces alterations of adjacent residues and may account for subunit asymmetry. (D) Alignment of the P3 structure to a previously determined model of the chemotaxis array in *E. coli* (PDB ID: 3JA6) indicates that the P3 domain lies within ∼15 Å of the receptors (when measuring from peptide back-bone). CheA and CheW in this model are shown in grey.

**Fig. S15.**
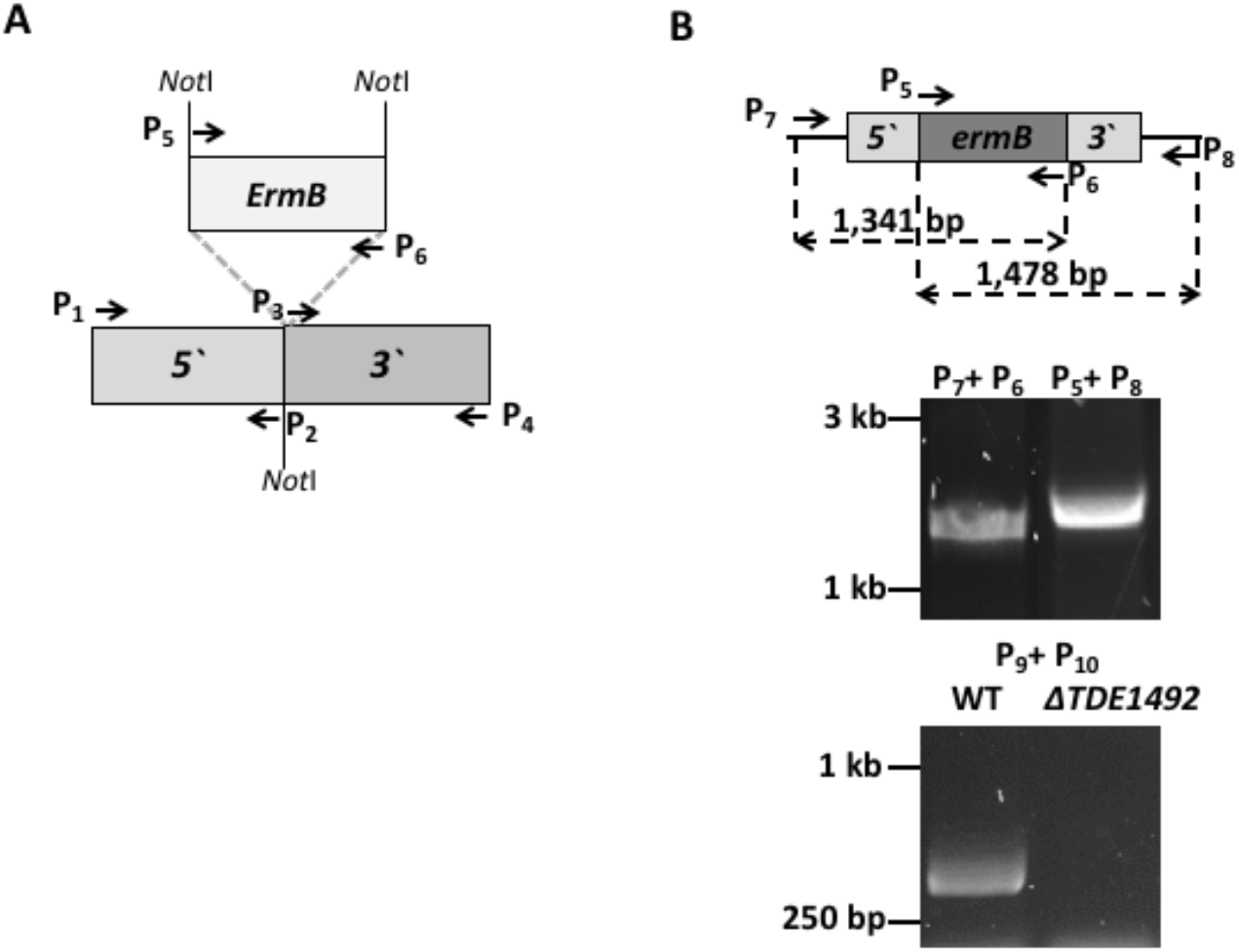
Diagrams illustrating construction of the *TDE1492::ermB* vector (A) for the targeted mutagenesis of *TDE1492 (781-1,308 nt)* by in-frame replacement of *TDE1492* using *ermB* cassette. These constructs were constructed by two-step PCR followed by DNA cloning. Arrows represent the relative positions and orientations of these primers, which are listed in Table S3. *ermB =* erythromycin resistance. (B) Characterization of the *ΔTDE1492* strain by PCR analysis. The top panel illustrates how the PCR analysis is designed; the bottom panel is the PCR results. Arrows represent the relative positions and orientations of these primers; the numbers are predicted sizes of PCR products generated by the corresponding primers. The primer P7 is located at the 5’-end of *TDE1492,* P_6_ at the 3’-end of *ermB,* P_5_ at the 5’-end of *ermB,* P_8_ at the flanking region of *TDE1492,* P_9_ at the middle of *TDE1492,* and P_10_ at the 3’-end of *TDE1492*. The sequences of these primers are listed in Table S3.

**Fig. S16.**
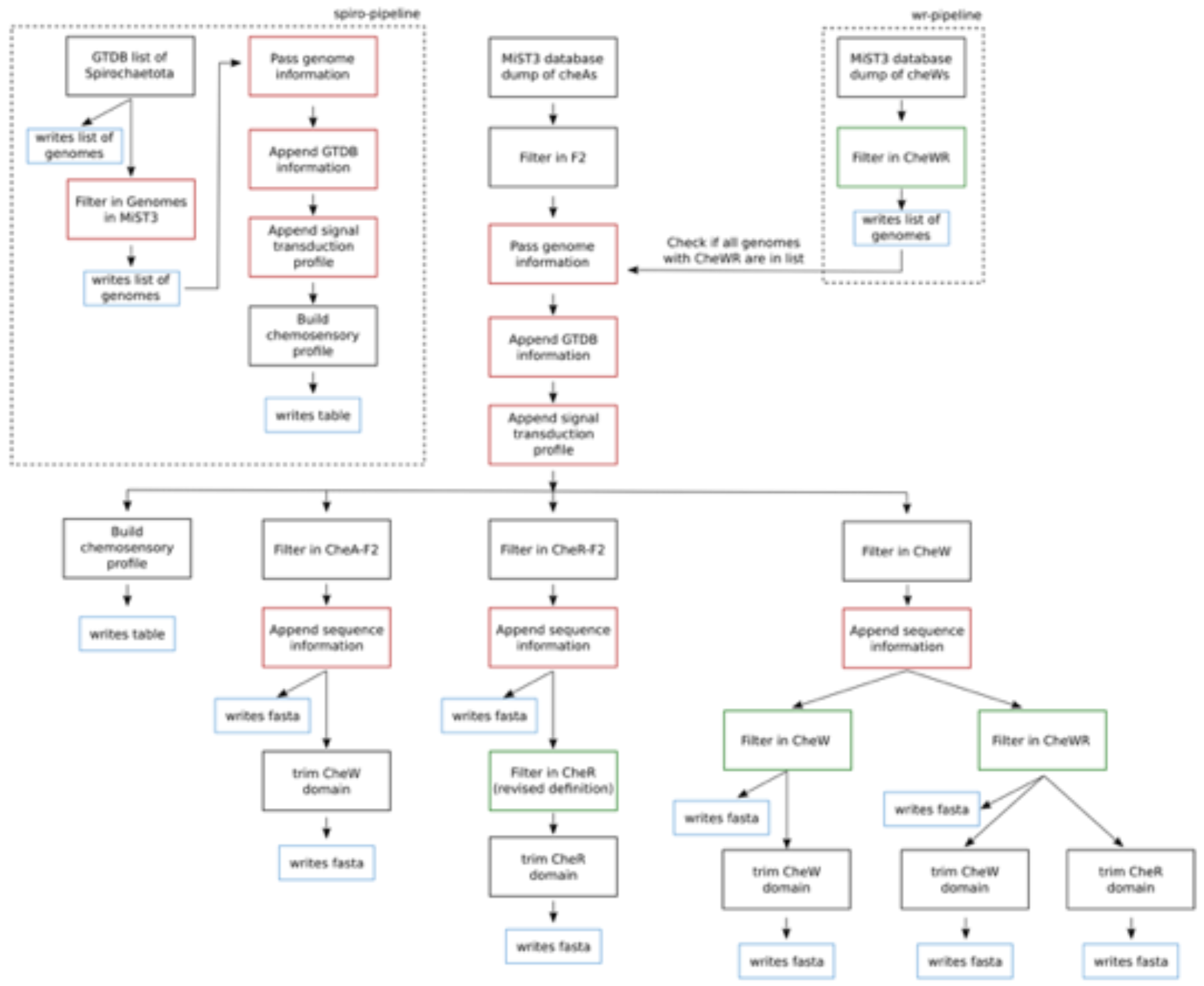
Flowchart of the three major pipelines used to produce the bioinformatics datasets. Steps marked in red represents fetching information from MiST3 database, in green are steps requiring RegArch as a filter, and in blue indicate writing data to file.

**Table S1A.** Statistics for the average angle between the *Td* cell axis and the ‘strands’ of CheA:CheW rings.

| *Td* strain | Average angle (°) |
| --- | --- |
| WT (n=9) | 9.67 +/- 9.21 |
| Δ2498 (n=9) | 9.55 +/- 8.75 |
| Δ2498Δ2496 (n=8) | 12.18 +/- 7.62 |
| **Combined average:** | **10.40 +/- 8.59** |

**Table S1B.**
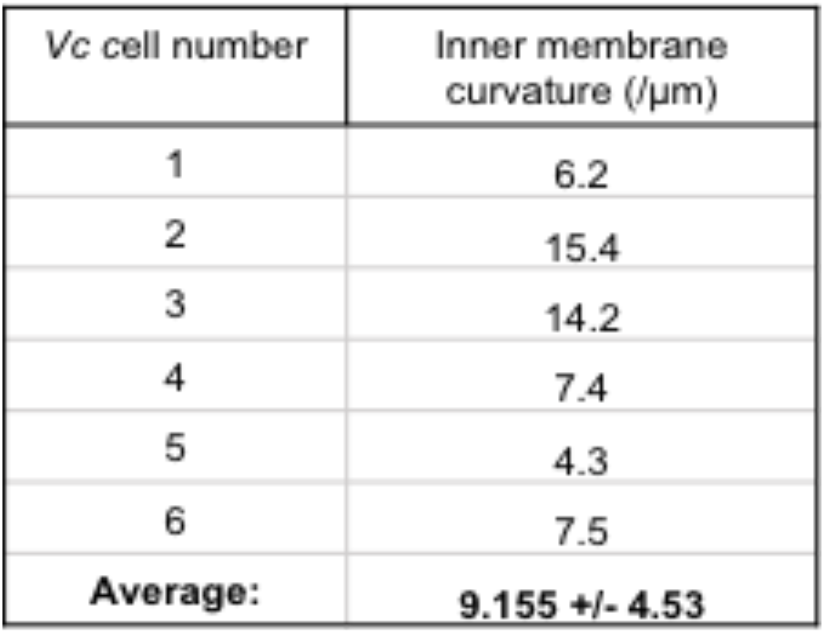
Statistics of inner membrane curvature for *6Vc* mini-cells.

**Table S1C.** Statistics of inner membrane and baseplate curvature for 10 WT *Td* cells.

| *Td* cell number | Inner membrane curvature (/μm) | Baseplate curvature (/μm) |
| --- | --- | --- |
| 1 | 35.0 | 49.7 |
| 2 | 25.4 | 54.0 |
| 3 | 34.2 | 63.7 |
| 4 | 43.1 | 96.1 |
| 5 | 30.8 | 99.7 |
| 6 | 41.1 | 55.7 |
| 7 | 39.6 | 74.6 |
| 8 | 33.1 | 47.7 |
| 9 | 46.2 | 47.8 |
| 10 | 29.1 | 67.1 |
| **Average:** | **35.8 +/- 6.6** | **65.6 +/- 19** |

**Table S2.**
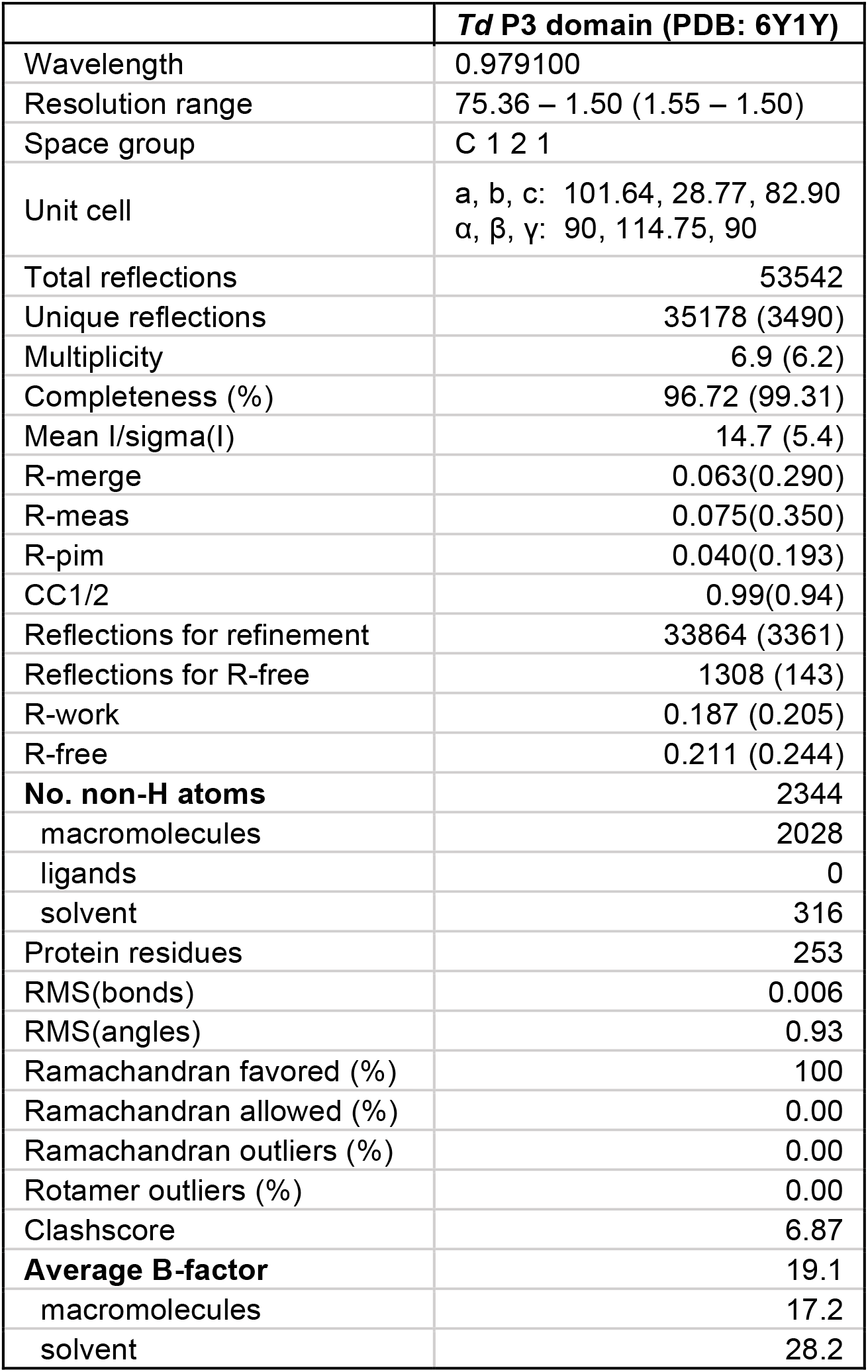
Data collection and refinement statistics for *Td* CheA P3 domain.

|  | ***Td* P3 domain (PDB: 6Y1Y)** |
| --- | --- |
| Wavelength | 0.979100 |
| Resolution range | 75.36 – 1.50 (1.55 – 1.50) |
| Space group | C 1 2 1 |
| Unit cell | a, b, c: 101.64, 28.77, 82.90<br>$\alpha$, $\beta$, $\gamma$: 90, 114.75, 90 |
| Total reflections | 53542 |
| Unique reflections | 35178 (3490) |
| Multiplicity | 6.9 (6.2) |
| Completeness (%) | 96.72 (99.31) |
| Mean I/sigma(I) | 14.7 (5.4) |
| R-merge | 0.063(0.290) |
| R-meas | 0.075(0.350) |
| R-pim | 0.040(0.193) |
| CC1/2 | 0.99(0.94) |
| Reflections for refinement | 33864 (3361) |
| Reflections for R-free | 1308 (143) |
| R-work | 0.187 (0.205) |
| R-free | 0.211 (0.244) |
| **No. non-H atoms** | 2344 |
| macromolecules | 2028 |
| ligands | 0 |
| solvent | 316 |
| Protein residues | 253 |
| RMS(bonds) | 0.006 |
| RMS(angles) | 0.93 |
| Ramachandran favored (%) | 100 |
| Ramachandran allowed (%) | 0.00 |
| Ramachandran outliers (%) | 0.00 |
| Rotamer outliers (%) | 0.00 |
| Clashscore | 6.87 |
| **Average B-factor** | 19.1 |
| macromolecules | 17.2 |
| solvent | 28.2 |

**Table S3.**
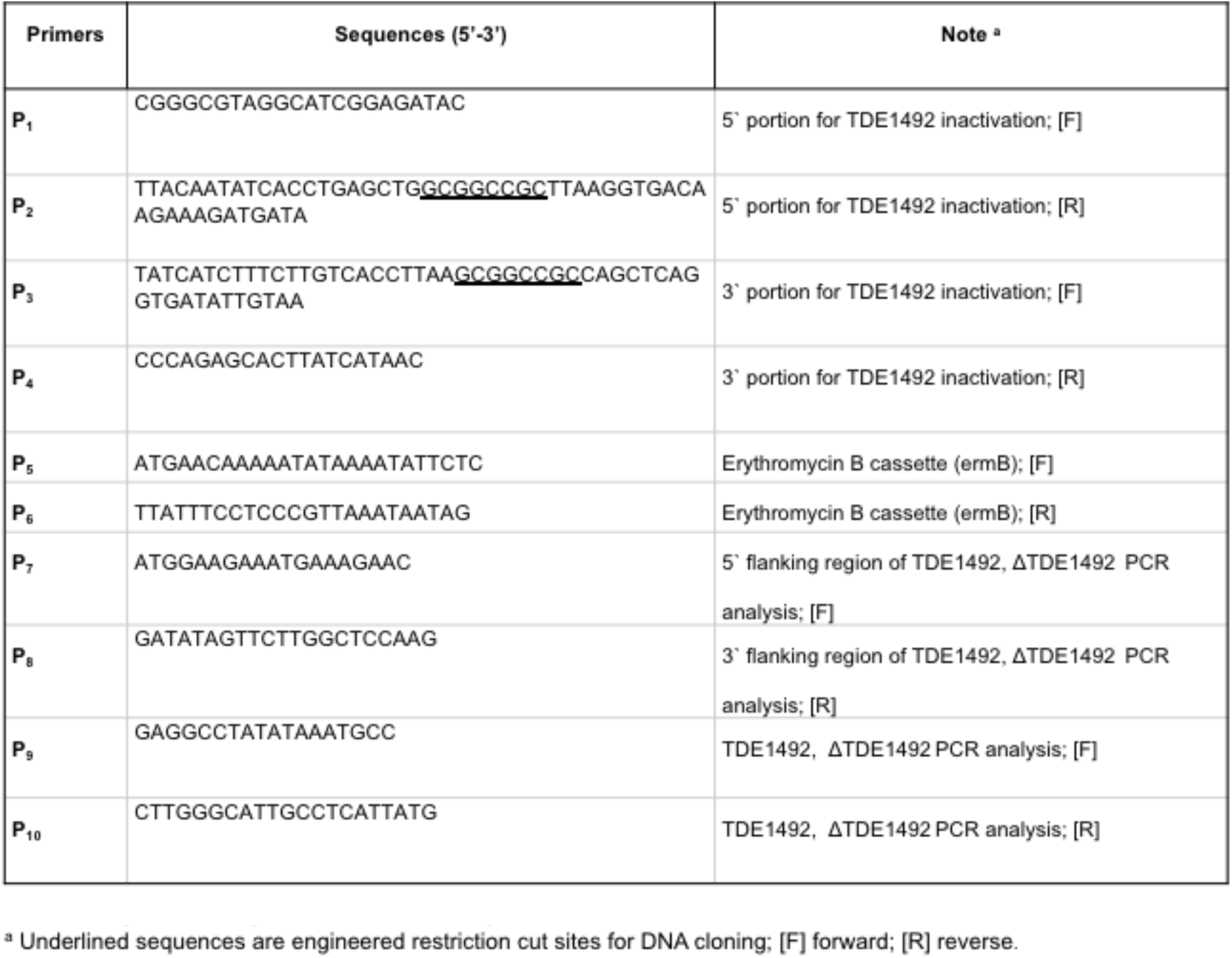
Oligonucleotide primers used in this study.

## Notes

#### Summary of Updates

The prokaryotic chemotaxis system is arguably the best-understood signaling pathway in biology, but most insights have been obtained from only a few model organisms and many studies have relied on artificial systems that alter membrane curvature1,2,3. In all previously described species, chemoreceptors organize with the histidine kinase (CheA) and coupling protein (CheW) into a hexagonal (P6 symmetry) extended array that is considered universal among archaea and bacteria4,5. Here, for the first time, we report an alternative symmetry (P2) of the chemotaxis apparatus that emerges from a strict linear organization of CheA in Treponema denticola cells, which possesses arrays with the highest native curvature investigated thus far. Using cryo-ET, we reveal that the Td chemoreceptor arrays assume a truly unusual arrangement of the supra-molecular protein assembly that has likely evolved to accommodate the high membrane curvature. The arrays have several additional atypical features, such as an extended dimerization domain of CheA and a variant CheW-CheR-like fusion protein that is critical for maintaining an ordered chemosensory apparatus in an extremely curved cell. Furthermore, the previously characterized Td oxygen sensor ODP influences array integrity and its loss substantially orders CheA. These results suggest a greater diversity of the chemotaxis signaling system than previously thought and demonstrate the importance of examining transmembrane systems in vivo to retain native membrane curvature.

